# Functional characterization of thousands of type 2 diabetes-associated and chromatin-modulating variants under steady state and endoplasmic reticulum stress

**DOI:** 10.1101/2020.02.12.939348

**Authors:** Shubham Khetan, Susan Kales, Romy Kursawe, Alexandria Jillette, Steven K. Reilly, Duygu Ucar, Ryan Tewhey, Michael L. Stitzel

## Abstract

A major goal in functional genomics and complex disease genetics is to identify functional *cis-*regulatory elements (CREs) and single nucleotide polymorphisms (SNPs) altering CRE activity in disease-relevant cell types and environmental conditions. We tested >13,000 sequences containing each allele of 6,628 SNPs associated with altered *in vivo* chromatin accessibility in human islets and/or type 2 diabetes risk (T2D GWAS SNPs) for transcriptional activity in ß cell under steady state and endoplasmic reticulum (ER) stress conditions using the massively parallel reporter assay (MPRA). Approximately 30% (n=1,983) of putative CREs were active in at least one condition. SNP allelic effects on *in vitro* MPRA activity strongly correlated with their effects on *in vivo* islet chromatin accessibility (Pearson r=0.52), i.e., alleles associated with increased chromatin accessibility exhibited higher MPRA activity. Importantly, MPRA identified 220/2500 T2D GWAS SNPs, representing 104 distinct association signals, that significantly altered transcriptional activity in ß cells. This study has thus identified functional ß cell transcription-activating sequences with *in vivo* relevance, uncovered regulatory features that modulate transcriptional activity in ß cells under steady state and ER stress conditions, and substantially expanded the set of putative functional variants that modulate transcriptional activity in ß cells from thousands of genetically-linked T2D GWAS SNPs.

## Introduction

Studies over the past decade have identified millions of single nucleotide polymorphisms (SNPs) in the human population^1^. Genome-wide association studies (GWAS) have linked thousands of these SNPs to variability in physiological traits and disease susceptibility^2^, including type 2 diabetes (T2D)^3^. The overwhelming majority of GWAS SNPs reside in non-coding, regulatory regions of the genome^3, 4^, implicating altered transcriptional regulation as a common molecular mechanism underlying disease risk. Compared to protein-coding regions, predicting regulatory functions of non-coding sequences, and the effect of SNPs within them, remains a significant challenge.

Approaches such as ATAC-seq identify regions of accessible chromatin as a marker of *in vivo* cis-regulatory elements (CREs)^5^. However, CREs identified by chromatin accessibility (ATAC-seq peaks) vary significantly in length. Moreover, ATAC-seq peaks are characterized by 1 or more summits, which have higher chromatin accessibility than flanking genomic regions also within ATAC-seq peaks^6^. Few studies have explored the relationship between transcriptional activity and open chromatin regions defined by ATAC-seq^7, 8^. For instance, it is not known whether higher chromatin accessibility within ATAC-seq peaks also corresponds to enhanced transcriptional activity, or whether proximity to ATAC-seq peak summits influences the likelihood for a SNP to affect enhancer activity. Studies designed to identify chromatin accessibility quantitative trait loci (caQTL) are being increasingly applied to nominate SNPs altering CRE activity^9, 10^. Previously, we identified 2,949 caQTLs in human islets^11^, which were significantly enriched for type 2 diabetes (T2D)-associated SNPs, highlighting the utility of this approach to nominate putative disease-associated SNPs affecting CRE activity in relevant cell types. However, caQTLs are correlative associations, and high throughput assays to directly test variant effects are needed to experimentally establish causality.

Massively parallel reporter assays (MPRAs) have been developed as a functional genomics platform to interrogate the enhancer potential of thousands of sequences simultaneously^12^ under a variety of (patho)physiologic conditions. By introducing nucleotide changes in a given sequence of interest, the effect of naturally occurring sequence variants in the human population on MPRA activity can also be elucidated^13^. Several studies have employed MPRA to identify functional SNPs associated with red blood cell traits^14^, adiposity^15^, osteoarthritis^16^, cancer and eQTLs^13^.

Although studies over the past several years implicate altered islet CRE function in T2D genetic risk and progression, they have colocalized only ∼1/4 of T2D-associated loci to altered chromatin accessibility and/or gene expression levels in islets^11, 17–20^. We hypothesize that this is partially because previous studies measured chromatin accessibility (caQTLs) and gene expression (eQTL) in islets cultured under steady state conditions^11, 21^, consequently missing important gene-environment interactions characteristic of a complex disorder like T2D. For example, endoplasmic reticulum (ER) stress, which is elicited by the high demands of insulin production and secretion on protein folding capacity in β cells, leads to (patho)physiologic adaptations. Low levels of ER stress are a signal for β cells to proliferate as a mechanism to meet higher insulin secretion demands^22^. However, uncompensated ER stress induced by genetic and/or environmental perturbations leads to β cell failure and T2D^23^.

Here, we applied MPRA to directly test >13,000 sequences, including those overlapping T2D GWAS and islet caQTL SNPs, for their ability to activate ß cell transcription in steady state conditions and after exposure to the ER stress-inducing agent thapsigargin or DMSO solvent control. We identified motifs and features of sequences affecting MPRA activity in each condition and linked allelic effects on MPRA activity to T2D genetics and *in vivo* islet chromatin accessibility. These results demonstrate the power of MPRA to elucidate *in vivo* relevant ß cell transcriptional activating sequences and define the regulatory grammar of these sequences under steady state and stress-responsive conditions. We nominate 54 putative causal variants among multiple genetically-linked islet dysfunction and T2D GWAS SNPs for further study.

## Results

### Selection and testing of sequences for MPRA activity in β cells

To test putative islet β cell regulatory sequences for their ability to enhance transcription from a minimal promoter and to identify SNPs that alter regulatory activity, we generated an MPRA library containing: i) 1,910 SNPs significantly associated with changes in human islet chromatin accessibility (caQTLs)^11^; ii) 2,218 control SNPs overlapping human islet ATAC-seq peaks, which were reported to have no correlation with changes in human islet chromatin accessibility^11^; and iii) 2,500 index and genetically linked (r^2^>0.8) SNPs/indels corresponding to 259 T2D association signals in the NHGRI/EBI GWAS Catalogue (Figure 1a, Table S2, and Methods). In total, 13,628 two-hundred base pair (bp) sequences from the human genome were included in this MPRA library.

**Figure 1.**
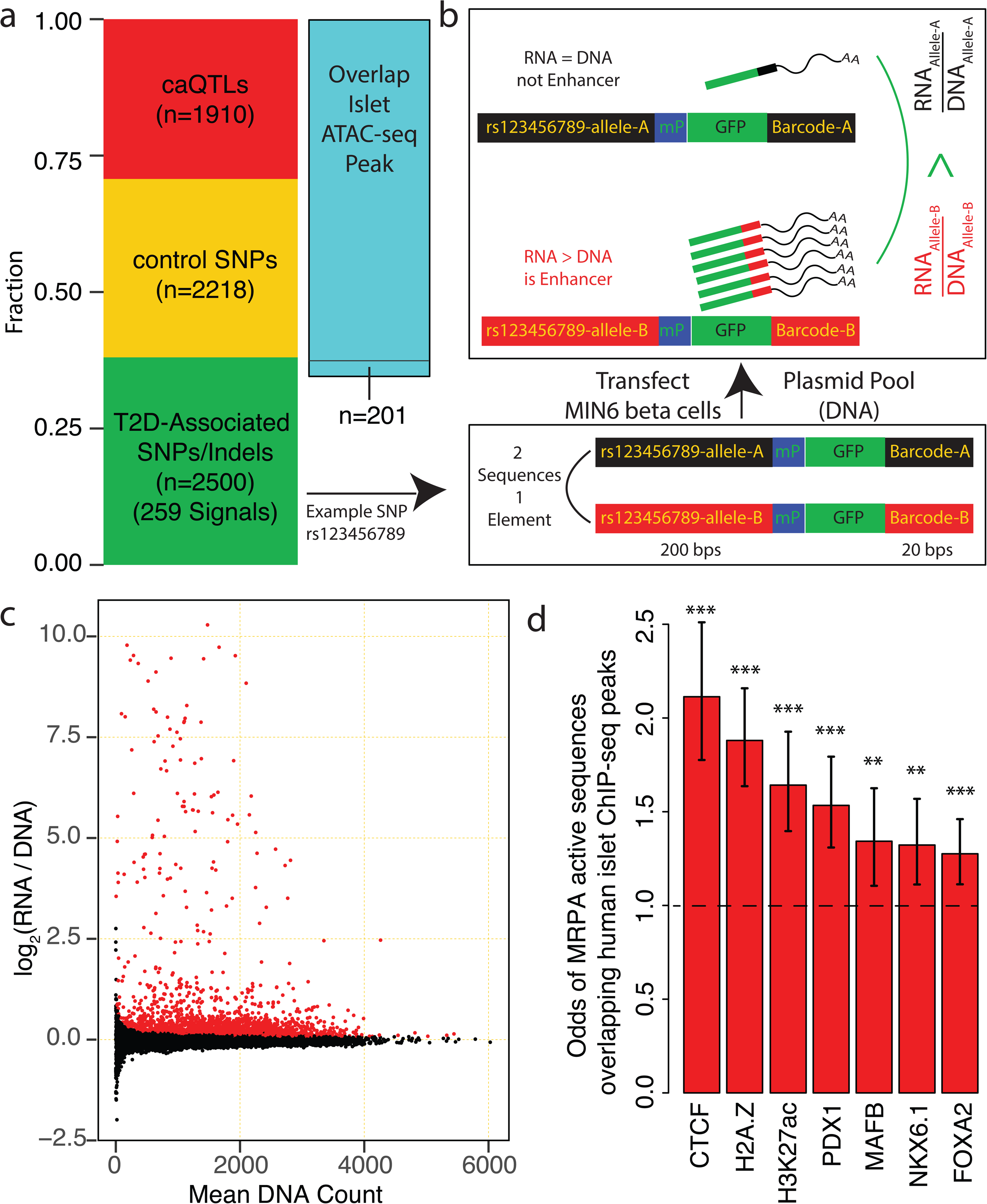
Massively Parallel Reporter Assay (MPRA) identifies beta cell transcription activating sequences. **(a)** Features of the 6,628 two hundred base pair (bp) elements selected for MPRA library construction and testing. 4,329 elements overlap islet ATAC-seq peaks, of which 1,910 contain SNPs associated with altered *in vivo* chromatin accessibility (caQTLs). 2,500 elements contain T2D-associated SNPs (index or linked (r^2^>0.8)) reported in the NHGRI/EBI GWAS catalog, 201 of which overlap islet ATAC-seq peaks. **(b)** Schematic of MPRA approach. A mock SNP, rs123456789, is used as an example to show how MPRA activity and allelic skew is measured for each 200 bp regulatory element. mP = minimal promoter. **(c)** Log_2_ fold-change in barcode/sequence counts in plasmid input (DNA) and *gfp* mRNA transcripts (RNA) 30 hr after transfection of MPRA library into MIN6 cells grown in standard culture conditions (DMEM, 25 mM glucose). Red points indicate sequences with MPRA activity (FDR<1%; n = 2224/13,628). **(d)** Barplot showing the odds of MPRA active sequences overlapping *in vivo* human islet TF ChIP-seq^17^ peaks compared to MPRA-inactive sequences. ***p<0.001; **p<0.01, Fisher’s Exact Test.

MIN6 mouse β cells have been extensively employed to study human islet regulatory sequences because human and mouse transcription factor (TF) expression and DNA binding motifs are extensively conserved. MIN6 cells express many β cell-specific TFs, such as *Pdx1*, *Nkx6.1* and *Foxa2*, and in a comparison with nine human tissues/cell-types, the chromatin accessibility profile of MIN6 mouse β cells most resembled that of human islets (Supplementary Figure 1). To test human sequences for islet β cell transcriptional activity, we transfected the MPRA plasmid library into five independent batches of MIN6 β cells under standard culture conditions (25mM glucose, Methods). Thirty hours later, transfected cells were harvested for RNA isolation, *GFP* mRNA capture, and Illumina sequencing of the regulatory sequence-associated barcodes (Figure 1b). As anticipated, principal component analysis indicated that RNA expression of the transfected MPRA libraries was highly correlated between the five biological replicates and clustered distinctly from the MPRA plasmid library input (Supplementary Figure 2, compare RNA vs. DNA). 2,224/13,628 sequences (16.3%), representing 1,372 distinct elements, exhibited significantly higher counts after transfection compared to the plasmid library input under standard culture conditions (FDR<1%; Figure 1c; Supplementary Table 2). We refer to these sequences as ‘MPRA active’ throughout the remainder of the manuscript. MPRA active sequences were significantly enriched for *in vivo* binding by human islet-specific TFs (PDX1, NKX6.1 and FOXA2; ChIP-seq; Figure 1d), suggesting that MPRA identifies human islet β cell regulatory sequences with *in vivo* relevance and validating MIN6 as a powerful cellular model to test human sequences for islet β cell transcriptional activity.

### Identification of β cell regulatory elements responding to ER stress

The high secretory burden of insulin production, processing, and secretion makes β cells particularly susceptible to ER stress. In fact, activation of the unfolded protein response (UPR) and ER stress has been implicated in the genetic etiology and pathophysiology of both monogenic^24, 25^ and type 2 diabetes^23^. Thapsigargin (TG) blocks calcium transport into the ER lumen and is widely used to induce ER stress in cells^26–33^. Therefore, to identify sequences whose ß cell transcriptional activity is modulated by ER stress, MIN6 cells transfected with the MPRA library were grown in standard culture media (25 mM glucose) supplemented with 250 nM TG or DMSO solvent control for 24 hours (Figure 2a). As expected, exposing MIN6 cells to 250 nM TG induced expression of ER stress response genes such as *Ddit3* (Chop), *Hspa5*, and *Edem1,* and reduced *Ins2* expression (Figure 2b). Compared to the DMSO solvent control, ER stress decreased MPRA activity of 656 sequences (mapping to 449 elements) and increased MPRA activity of 328 sequences (mapping to 275 elements), respectively, at FDR <1% (Figure 2c).

**Figure 2.**
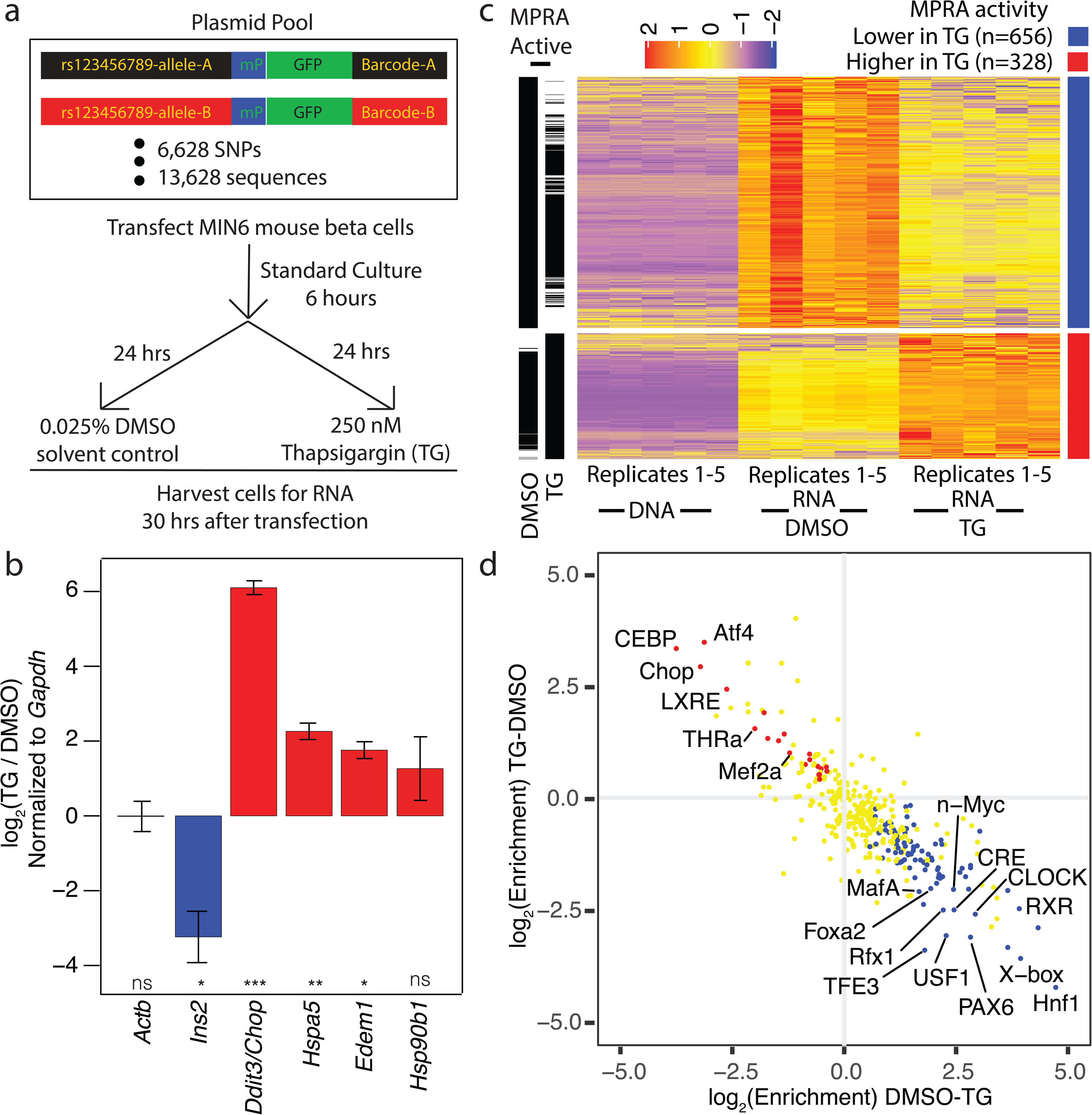
MPRA identifies endoplasmic reticulum (ER) stress-responsive transcriptional activating sequences. **(a)** Experimental design to identify ER stress-responsive transcription activating sequences. **(b)** RT-qPCR of ER stress response genes (*Ddit3, Hspa5, Edem1,* and *Hsp90b1)*, *Ins2,* and the control gene *Actb* in MIN6 cells exposed to the ER stress inducer thapsigargin (TG, 250 nM) or dimethylsulfoxide (DMSO) solvent control for 24 hours. Bar plots show mean +/-SEM from three biological replicates. *p<0.05; ** p<0.01; *** p<0.001, paired t-test. **(c)** Heatmap of sequences with higher (far right column, red bar; n=328, mapping to 275 elements) or lower MPRA activity (blue bar; n=656, mapping to 449 elements) under ER stress (FDR<1%). Red annotation bars to the left of the heatmap indicate sequences identified as MPRA active in DMSO and/or TG based on their sequence counts in RNA vs. plasmid DNA input. **(d)** Comparison of transcription factor (TF) binding motifs enriched in elements with higher (n=275) or lower (n=449) MPRA activity under ER stress. Red and blue dots denote TF motifs significantly enriched in elements with higher or lower MPRA activity in ER stressed MIN6 beta cells, respectively (FDR<1%). Yellow dots denote TF motifs with no significant enrichment in either comparison.

We compared these differentially active sequences to identify TF motifs that may be mediating their transcriptional activity differences. Elements enriched for sequence binding motifs of β cell-specific TFs that mediate insulin gene transcription and beta cell function^34, 35^, such as MAFA, FOXA2 and PAX6, had lower MPRA activity under ER stress (Figure 2d). Concordantly, we observed a significant decrease in insulin gene expression (Figure 2b), suggesting ER stress leads to inactivation of these β cell-specific TFs. It is known that uncompensated ER stress leads to preferential ATF4 translation and induction of *DDIT3/*CHOP^36, 37^. Consistently, we found that motifs for these TFs were enriched in elements with higher MPRA activity after ER stress induction (Figure 2d). Activity measurements with MPRA, therefore, recapitulate TF dynamics during ER stress and identify β cell regulatory elements that respond to ER stress.

### ATAC-seq peak summits are enriched for TF binding motifs and MPRA activity

In total, 30% of sequences containing one or both SNP alleles were MPRA active in at least one treatment condition (Figure 3a, n=1983/6628, FDR ≤1%). MPRA active sequences were significantly enriched for 114 TF binding motifs (Supplementary Table 3), including those of islet TFs such as HNF1, MAFA, RFX5, and FOXA2. Human-mouse sequence similarity did not influence the probability that a tested human sequence was MPRA active in MIN6 (black dashed line in Figure 3b; Supplementary Figure 3). As anticipated, sequences overlapping regions with *in vivo* evidence of transcriptional regulatory potential in human islets (ATAC-seq peaks) were significantly more likely to be identified as MPRA active regardless of the sequence similarity between human and mouse (Figure 3b).

**Figure 3.**
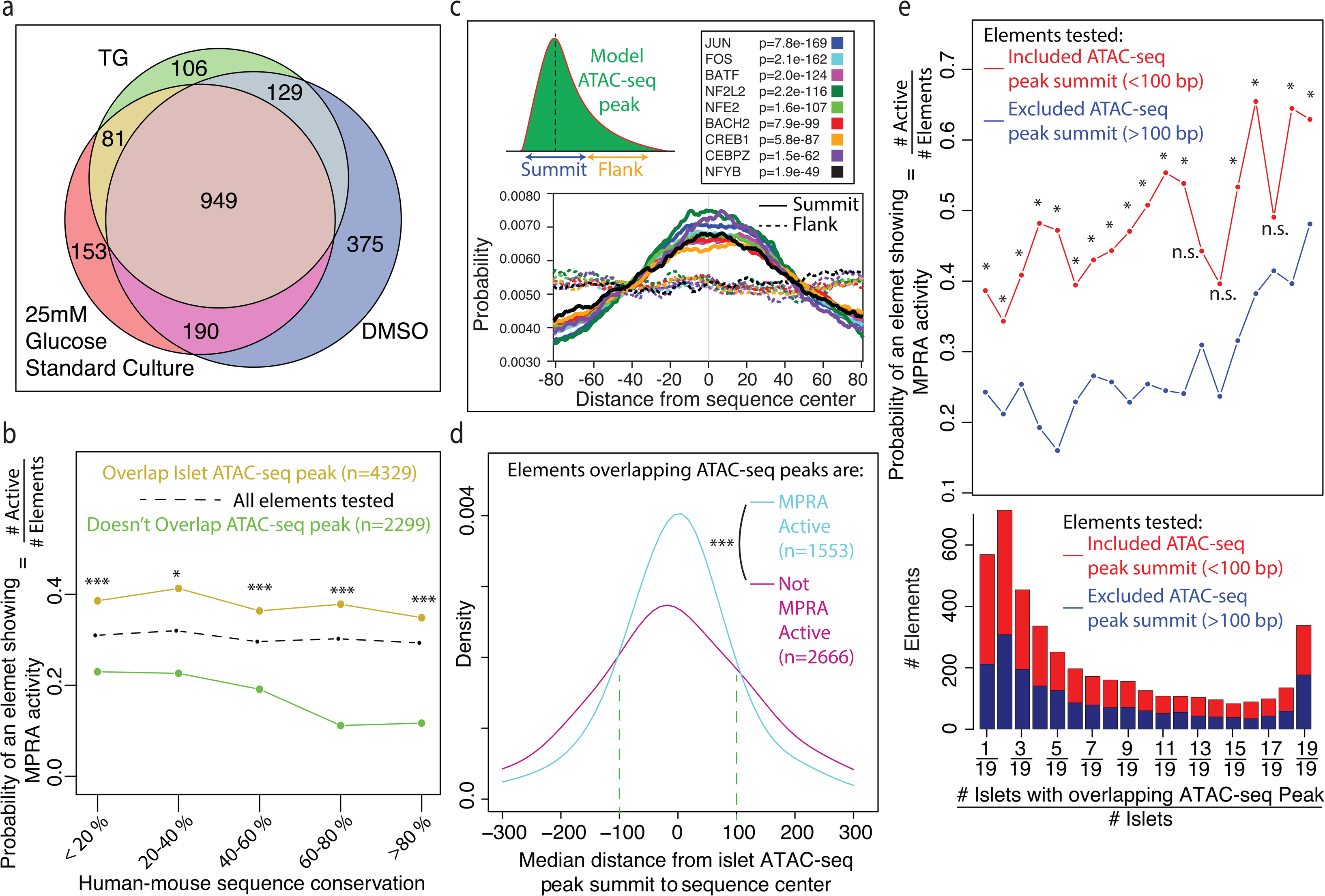
ATAC-seq peak summits are enriched for MPRA activity. **(a)** Venn diagram showing the number of elements with at least one allele identified as MPRA active in standard culture, TG, and DMSO conditions (FDR<1%). Note that elements identified as MPRA active in ≥1 experimental condition may still exhibit quantitative differences in MPRA activity under ER stress. **(b)** The probability that an element has significant MPRA activity (y-axis; at least 1 allele; any experimental condition) is shown as a function of sequence similarity (x-axis) between human and mouse genomes (expected probability is the black dotted line). Elements overlapping islet ATAC-seq peaks (brown line) had a significantly higher probability of MPRA activity than elements that did not overlap ATAC-seq peaks (green line), regardless of human-mouse sequence similarity. ***, **, and * indicate Bonferroni-corrected Fisher’s Exact Test p<0.001, 0.01, 0.05, respectively. **(c)** 200 bp genomic regions around human islet ATAC-seq peak summits were compared to 200 bp regions also within ATAC-seq peaks, but flanking the peak summit. ATAC-seq peak summits were significantly enriched for many TF motifs compared to genomic regions that overlap ATAC-seq peaks but are further away from the summit. **(d)** Elements where the SNP (center of all tested loci) was ± 100 bp (green dashed lines) from ATAC-seq peak summit were more likely to be identified as MPRA active. This effect is restricted to SNPs within 100 bps of the ATAC-seq peak summit because our MPRA library was designed to test only the 200 bp sequence on either side of a given SNP of interest. ATAC-seq peak summits were therefore included in the sequences tested only if the distance of the SNP was within 100 bps of the ATAC-seq peak summit. **(e)** (Top) The probability of an element (at least one allele) being identified as MPRA active (y-axis) binned by the number of islet donors (out of 19 total) in whom the ATAC-seq peak was present (x-axis). Elements where the ATAC-seq peak summit was included in the sequence tested, i.e., distance of SNP to ATAC-seq peak summit was less than 100 bps (red), had a significantly higher probability of being identified as MPRA active. * indicates FDR < 10%, Fisher’s Exact Test. (Bottom) Stacked barplot of the number of times an ATAC-seq peak overlapping the MPRA sequence tested is detected in the cohort (n = 19 islet donors).

Notably, the majority (60%) of elements accessible in human islets were not MPRA active (Figure 3b). Islet ATAC-seq peaks vary in length (100-3500 bps). Moreover, within an ATAC-seq peak, there are differences in chromatin accessibility levels at nucleotide resolution, where peak summits are locations with higher read counts than the flanking regions (schematic in Figure 3c; example in Figure 5a). Peak summits are significantly enriched in TF binding motifs compared to flanking regions within the same ATAC-seq peak (Figure 3c and Supplementary Figure 4). Therefore, for MPRA library sequences overlapping ATAC-seq peaks, we examined the distance of SNPs, which we designed to be in the center of all MPRA sequences tested, to the summit of ATAC-seq peaks in which they reside. Sequences closer to ATAC-seq peak summits were significantly more likely to be MPRA active (Figure 3d). However, this effect was observed only when SNPs were less than 100 bps from the ATAC-seq peak summit (dashed green line in Figure 3d). Since the MPRA library tests the 100 base pairs flanking both sides of a given SNP of interest, ATAC-seq peak summits were included in the MPRA sequences only if they were ≤100 bp from the SNP. Therefore, MPRA sequences overlapping ATAC-seq peaks were categorized based on whether the summit was within 100 bps of the SNP or not, i.e., whether the ATAC-seq peak summit was included in the sequence tested with MPRA or not. Sequences that included the ATAC-seq peak summit were indeed more likely to be active with MPRA, irrespective of the number of times a genomic region was accessible in a cohort of 19 islet donors (Figure 3e). Interestingly, inclusion of the ATAC-seq peak summit was predictive of MPRA activity even for genomic regions with accessibility in only one of 19 islet donors.

Together, these analyses indicate that sequences overlapping human islet ATAC-seq peaks were more likely to be active with MPRA in MIN6 mouse β cells than sequences outside peaks, regardless of sequence similarity between human and mouse. ATAC-seq peak summits are enriched for TF binding motifs, and exclusion of ATAC-seq peak summits among the elements overlapping ATAC-seq peaks, significantly decreased the probability of observing MPRA activity.

### SNPs proximal to ATAC-seq summits have a higher probability of altering MPRA activity and chromatin accessibility

To identify SNPs that modulate regulatory element activity, we assessed allelic differences (skew) in MPRA activity (schematic in Figure 1b) under each MIN6 beta cell culture condition (standard, 250nM TG, and DMSO solvent control). In total, 879 SNPs exhibited allelic skew at FDR <10% (Table S2; Supplementary Figure 5). Importantly, when allelic skew was detected in more than one experimental condition, the direction-of-effect was concordant for >98.5% of the SNPs (Supplementary Figure 5). Assessing allelic skew for SNPs across all conditions increased the power to detect allelic skew and identified both highly reproducible and condition-specific SNP allelic effects on MPRA activity.

SNPs associated with chromatin accessibility changes in islets (caQTLs) were significantly closer to the ATAC-seq peak summits than control SNPs, which are SNPs that reside in ATAC-seq peaks^11^ but were not associated with chromatin accessibility changes in islets (Figure 4a). Consequently, caQTL SNPs were more likely to be MPRA active than were control SNPs (Figure 4b). Among elements with MRPA activity, caQTL SNPs were also more likely to exhibit allelic skew than control SNPs (Figure 4c) and showed a significantly greater magnitude of skew between alleles (Figure 4d). Together, these results suggest that caQTL SNPs are closer to ATAC-seq peak summits and more likely to modulate the magnitude of both *in vitro* MPRA activity and *in vivo* chromatin accessibility, likely by disrupting TF binding motifs that are enriched in these regions.

**Figure 4.**
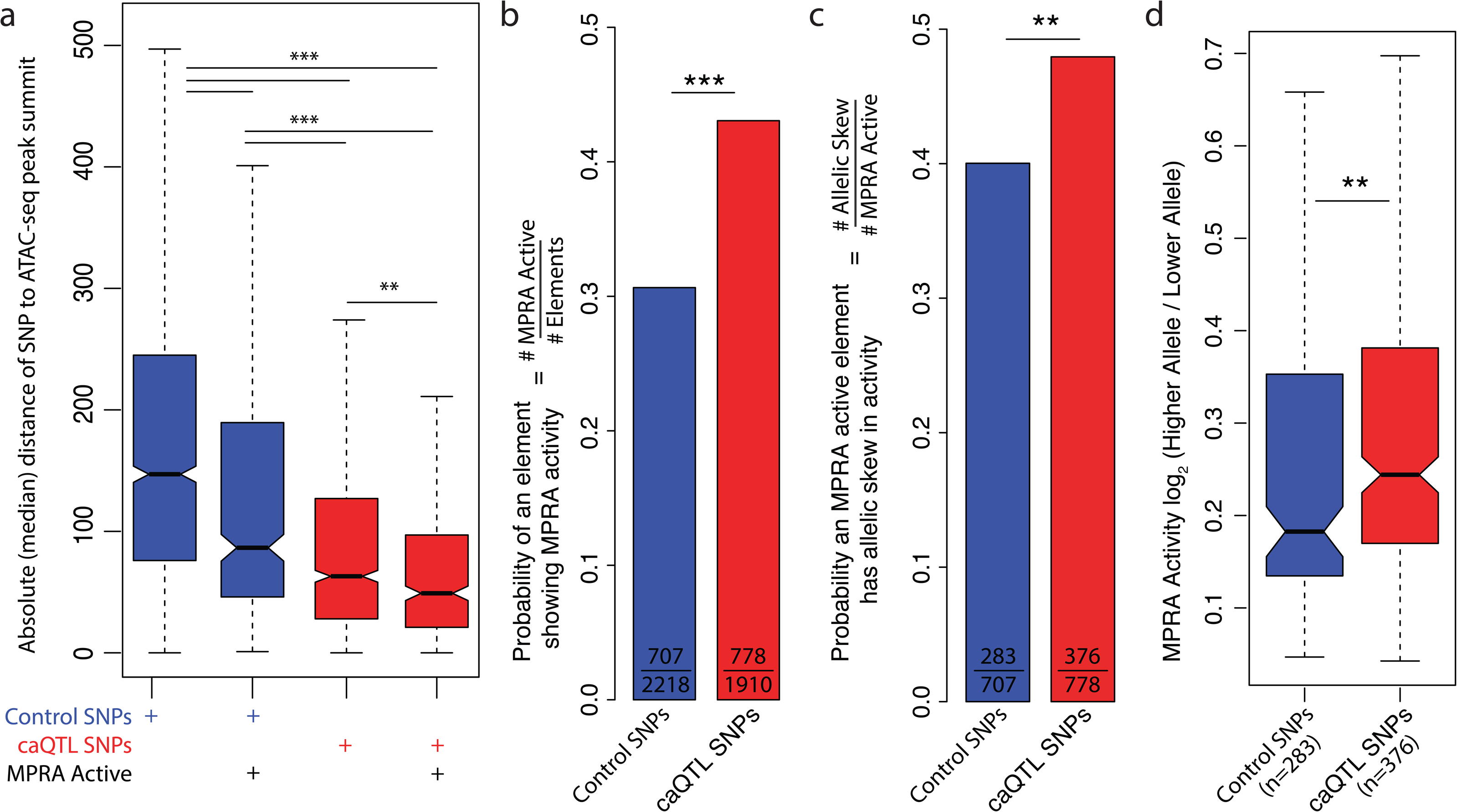
SNPs closer to ATAC-seq peak summits are more likely to affect MPRA activity and chromatin accessibility. **(a)** caQTL SNPs (blue bars) are closer to ATAC-seq peak summits than control SNPs (red bars), irrespective of MPRA activity (pairwise Wilcoxon tests with Bonferroni correction for multiple testing). **(b)** Elements containing caQTL SNPs are more likely to be identified as MPRA active (any experimental condition; Fisher’s Exact Test). **(c)** Among SNPs with at least 1 allele identified as MPRA active, caQTLs are significantly more likely show allelic skew (any experimental condition), suggesting polymorphisms proximal to ATAC-seq peak summits are more likely to affect MPRA activity (Fisher’s Exact Test). **(d)** Among SNPs with allelic skew in MPRA activity, caQTLs have a significantly larger difference in MPRA activity between alleles than control SNPs (Wilcoxon Test). For all panels, *, **, and *** indicate adjusted p <0.05, p < 0.01, and p < 0.001, respectively.

### Allelic effects on MIN6 beta cell MPRA activity and *in vivo* islet chromatin accessibility are significantly correlated

After identifying SNPs with allelic skew in MPRA activity, we investigated the relationship between SNP effects on *in vivo* islet chromatin accessibility (caQTLs) and *in vitro* MPRA activity. Figure 5a shows an example caQTL (rs17396537) for which islet donors with homozygous GG genotypes (n=6) exhibit higher chromatin accessibility than islet donors with GC (n=11) or CC (n=2) genotypes at this variant. Concordantly, the sequence containing the ‘G’ allele for rs17396537 exhibited higher MPRA activity than the one containing the ‘C’ allele (Figure 5a, b). 297/1910 lead caQTL SNPs demonstrated significant allelic skew in MPRA activity. Overall, caQTL and MPRA directions-of-effect for these 297 SNPs were highly correlated (Pearson R = 0.526, Figure 5b), and a significant concordance of direction was observed (i.e., alleles associated with higher chromatin accessibility also had higher MPRA activity) (n = 246/297; 82.8%).

**Figure 5.**
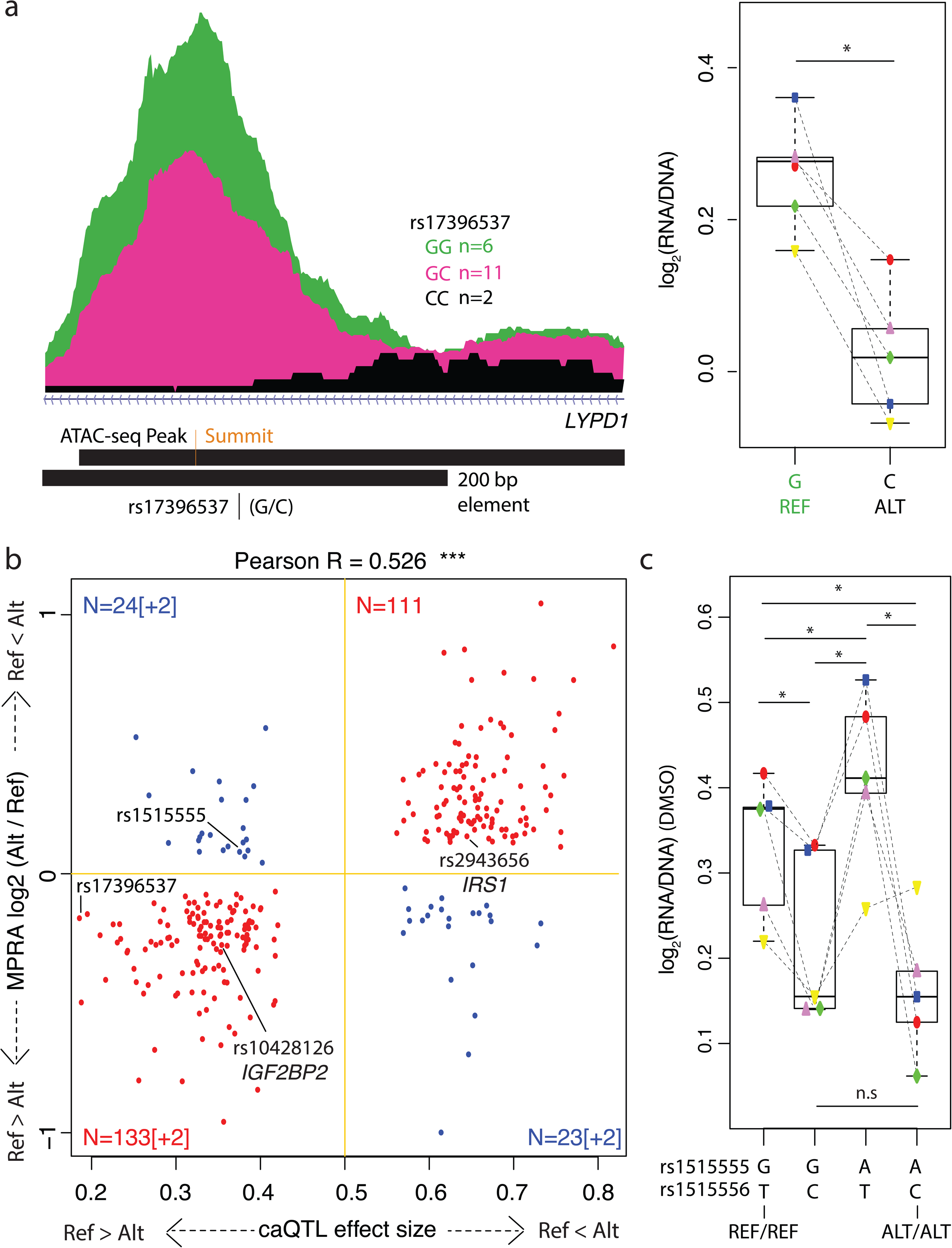
Alleles associated with higher chromatin accessibility also have higher MPRA activity. **(a)** (Left) caQTL example: Islet samples GG homozygous for rs17396537 (green; average ATAC-seq read counts) were associated with significantly higher chromatin accessibility than samples with GC (pink) or CC (black) genotypes (hg19 coordinates - chr2:133408787-133409072). (Right): Sequence with the reference ‘G’ allele at rs17396537 displayed significantly higher MPRA activity than the alternate ‘C’ allele. **(b)** Correlations between allelic effects on *in vivo* islet chromatin accessibility (x-axis) and MPRA activity (y-axis) for sequences containing islet caQTL SNPs. Quadrants 1 and 3 (red) correspond to SNPs where the allele associated with higher chromatin accessibility also had higher MPRA activity (concordant). The number of SNPs in each quadrant is indicated. Pearson R = 0.526, p value < 2.2e-16) among the caQTL SNPs with significant allelic skew in MPRA activity (FDR<10%). Brackets indicate points beyond the axis limits. For example, there are a total of 135 SNPs in the bottom left quadrant, 2 of which are beyond the axis limits. REF=hg19 reference allele; ALT=alternate allele. **(c)** As shown in panel b, the lead caQTL SNP rs1515555 demonstrated apparently discordant allelic effects between chromatin accessibility and MPRA activity. Since there is another SNP, rs1515556, 24 bps from the lead caQTL SNP (r^2^=0.99), sequences containing all four allelic combinations were tested in the MPRA library. The alternate allele of rs1515555 increased MPRA activity, whereas that of rs1515556 decreased MPRA activity. Ref/Ref for rs1515555 and rs1515556 exhibited higher MPRA activity than the Alt/Alt combination, which is concordant with its higher chromatin accessibility associated with Ref/Ref for rs1515555 and rs1515556.

A subset of lead caQTL SNPs (17.2%) showed discordant allelic effects on chromatin accessibility levels and MPRA activity (blue; Figure 5B). We hypothesized that discordance may be driven by a nearby SNP that has MPRA activity antagonistic to the lead caQTL SNP^10^. For example, the reference allele of the lead caQTL SNP, rs1515555, is associated with *in vivo* higher chromatin accessibility but lower MPRA activity (Figure 5b). This region contains a second SNP, rs1515556, 24 bps away from rs1515555 and in tight LD (r^2^ = 0.99). We therefore assessed sequences containing each of the four possible allelic combinations of these SNPs in the MPRA library for additive, antagonistic, or synergistic transcriptional effects. In comparison to rs1515555:rs1515556 REF:REF sequence (G:T in Figure 5c), the ALT:REF (A:T) sequence had higher MPRA activity, while the REF:ALT (G:C) sequence had lower MPRA activity (G:C). However, due to high LD between the 2 SNPs, both of these combinations (‘A:T’ and ‘G:C’; ALT:REF and REF:ALT; Figure 5c) are rarely observed in the human population. Sequences with either REF:REF (G:T) or ALT:ALT (A:C) at rs1515555:rs1515556 are more likely to be observed in individuals in the population. Comparison of these haplotypes revealed concordant effects on both chromatin accessibility and MPRA activity and underscores the importance of accounting for haplotypes when characterizing allelic effects.

For 94 lead caQTL SNPs, we tested for potential interactions with neighboring SNPs by including all 4 allelic combinations in the MPRA library. However, these were too few to clearly implicate antagonism between neighboring SNPs for the observed discordance between chromatin accessibility and MPRA activity. Therefore, we cannot rule out discordant SNPs as merely being false positives or mechanistic difference due to technical constraints of the assay. However, MPRA provides a unique opportunity to test and dissect the contributions of neighboring SNP to *in vivo* effects.

Allelic effects on MPRA activity and chromatin accessibility are therefore significantly correlated. Importantly, these results demonstrate that that MPRA detects *in vivo* relevant allelic effects on ß cell transcriptional activity.

### Functional identification of T2D SNPs altering β cell regulatory element activity

A key challenge to translate GWAS associations into a mechanistic understanding of the genes and pathways altered by T2D risk variants is identifying the SNP(s) that affect functional or active CREs among tens to hundreds of genetically linked, associated SNPs in each locus. The ability to functionally annotate groups of SNPs with empirical measurements of CRE modulation would help prioritize variants and expedite our ability to identify T2D causal alleles. To uncover transcriptional effects of T2D-associated SNP alleles, we tested sequences containing each allele of 2,500 index and genetically linked (r^2^>0.8) SNPs/indels for 259 association signals for T2D and related quantitative traits reported in the NHGRI/EBI GWAS Catalog. One or both alleles of 492 SNPs/indels were active by MPRA. Active sequences were enriched for the binding motifs of TFs that play important roles in β cell maturation and insulin secretion (Figure 6a), such as liver x receptor (LXR)^38, 39^, thyroid hormone receptor (THRa and THRb)^40, 41^, retinoic acid receptor (RARa)^42^, and BCL11A^43^. Allelic effects on MPRA activity were observed for approximately half of these elements (n=220/492), corresponding to 104 distinct T2D-associated loci (Supplementary Table 4). Importantly, this list included T2D-associated SNPs that were previously identified for their *in vivo* effects on islet chromatin accessibility and/or *in vitro* effects on luciferase reporter assays, such as rs7903146 (*TCF7L2*)^44, 45^, rs1635852 (*JAZF1*)^46^, rs10428126 (*IGF2BP2*)^11, 47^, and rs12189774 (*VEGFA*)^48^ (Figure 6b, Table 1). Importantly, MPRA has substantially expanded the set of T2D-associated GWAS SNPs with empiric effects on transcriptional activation in ß cells, and therefore nominated them for the first time as the putative causal/functional variants in these T2D loci. These include rs2881632 (*SDHAF4*), rs13096599 (*KBTBD8*), rs7783500 (*GCC1*), rs72697237 (*NOTCH2*), rs13405776 (*THADA*), rs17748864 (*PEX5L*), rs7957197 (*OASL*), and rs13026123 (*SRBD1*), which are highlighted in Table 1.

**Figure 6.**
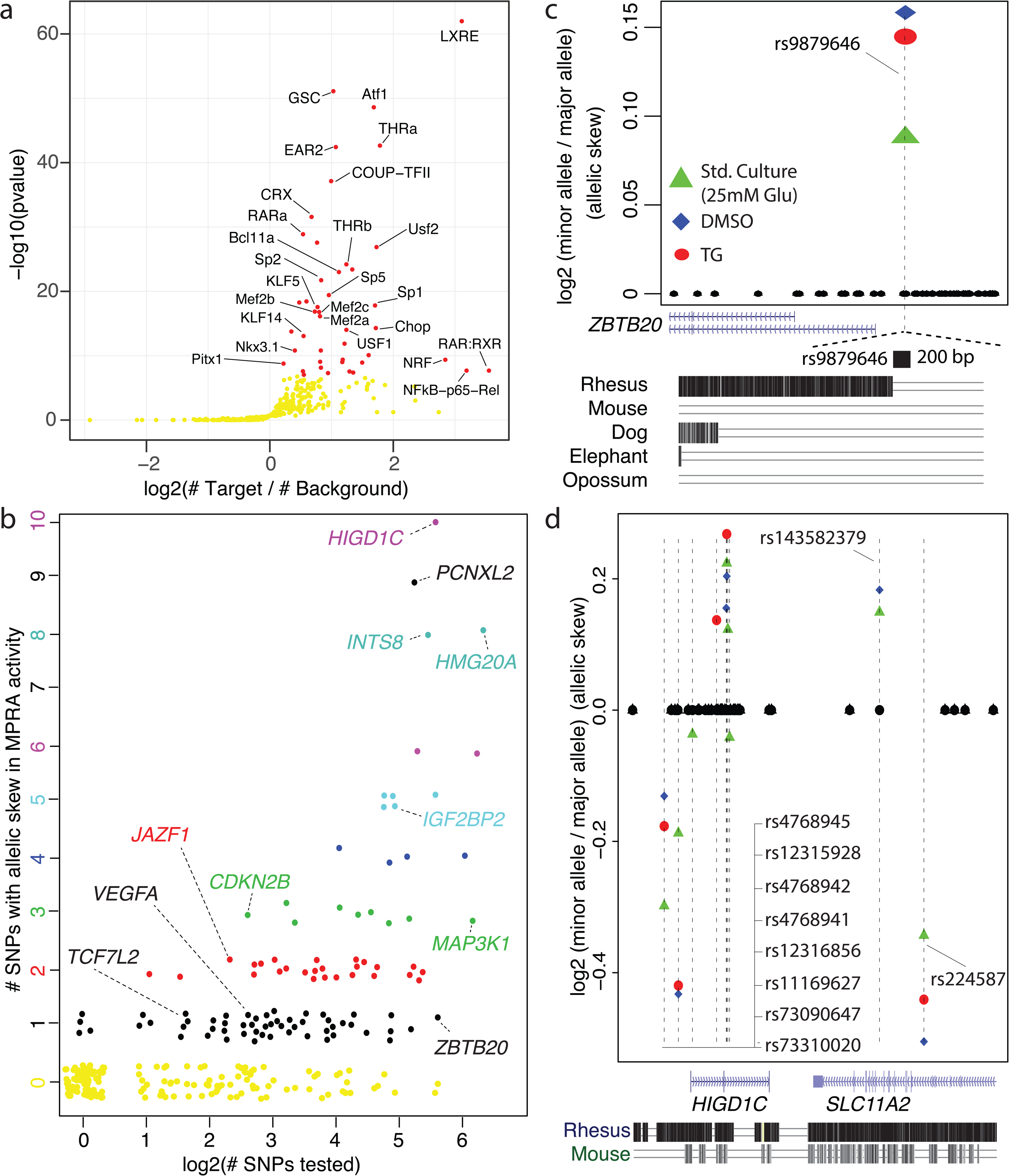
Functional identification of T2D SNPs altering β cell regulatory element activity. **(a)** TF binding motifs enriched at 492 elements containing T2D-associated SNPs with significant MPRA activity (at least one allele). Red dots denote significantly enriched TF binding motifs (FDR<1%). **(b)** MPRA identifies a subset of T2D-associated index and genetically linked (r^2^≥0.8) SNPs representing 259 signals from the NHGRI/EBI GWAS catalog (x-axis; log scale) that demonstrate significant allelic effects on MPRA activity (y-axis) in one or more condition tested. A total of 220 SNPs in linkage disequilibrium with 104 T2D-associated index SNPs show significant allelic effects on MPRA activity (FDR<10%). Jitter in the plot is used to visually separate individual points. **(c)** (Top). In total, 49 SNPs in LD (r^2^ > 0.8) with the T2D-associated index SNP rs73230612 were tested with MPRA (ZBTB20 locus). Allelic skew in MPRA activity was detected in the same direction across all 3 experimental conditions (standard culture, DMSO and TG) at 1 SNP only, rs987964. (Bottom; zoom-in) The 200 bp genomic region spanning rs987964 is specific to humans. **(d)** In total, 45 SNPs in LD (r^2^ > 0.8) with the T2D-associated index SNP rs12304921 were tested with MPRA (HIGD1C locus). Allelic skew in MPRA activity was detected at 10 SNPs.

For 54/104 T2D-associated signals, one SNP among those tested exhibited allelic effects on MPRA activity (black points in Figure 6b). For example, rs987964 was the only SNP of 49 tested in the *ZBTB20* locus (Figure 6c; Supplementary Table 5) that exhibited allelic effects on MPRA activity. For this and 53 additional T2D-associated signals, MPRA thus provides empiric evidence that the variant directly modulates CRE activity and assists in prioritizing variants for additional investigation as putative causal alleles (Supplementary Table 5). The number of SNPs with allelic skew was found to generally correlate with the number of SNPs tested per locus (Figure 6b). For example, 10/45 tested SNPs in LD with the index SNP rs12304921 in the *HIGD1C* locus had an allelic skew in MPRA activity (Figure 6d), not all of which were in the same direction with respect to the T2D risk alleles. Investigating the relative importance and potential interactions between these ten putative causal variants will be important to establish causality for this locus. These results underscore the value of assessing MPRA activity in disease-relevant cell types as a tool to nominate putative causal T2D SNPs based on systematic testing of their transcription-modulating effects in beta cells.

### ER stress modulates MPRA activity of T2D SNP-containing sequences

MPRA activity of some putative CREs were exclusive to one out of the three conditions tested. For example, ∼5% of those tested were identified as MPRA active exclusively in ER stressed beta cells (Figure 3a, n=106). However, we found that ER stress elicited substantial quantitative changes in MPRA activity of 724 CREs (Figure 2c). To investigate both the impact of ER stress on MPRA activity and how SNPs modify this response, we examined elements meeting two criteria: 1) allelic effects on MPRA activity was observed in at least one condition; and 2) a minimum of one allele showed differences in MPRA activity under ER stress (Figure 7a).

**Figure 7.**
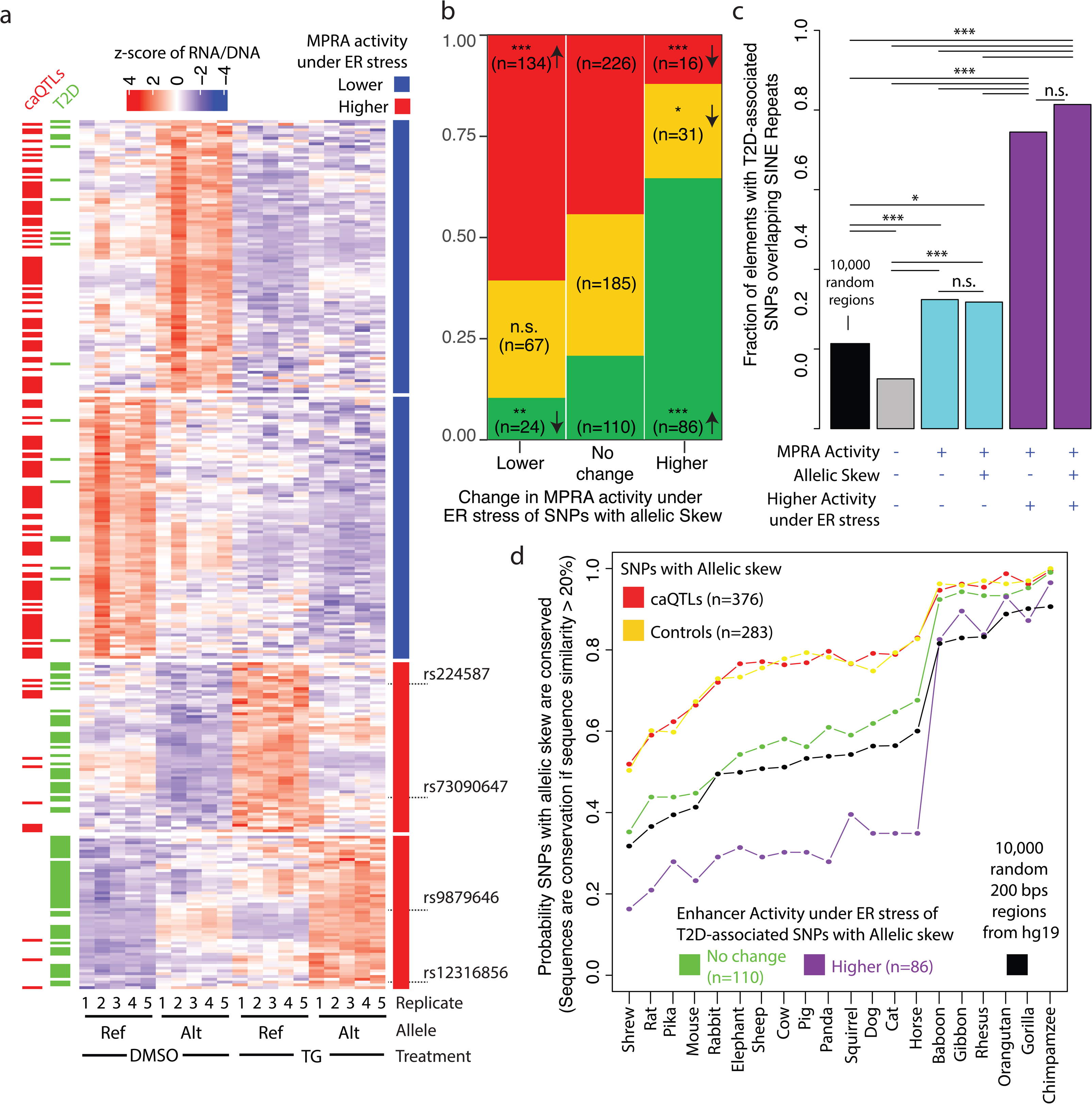
ER stress increases MPRA activity of 40% of T2D-associated SNP containing sequences with allelic skew. **(a)** Among the SNPs with allelic skew (any condition), the heatmap includes those with at least 1 allele showing significantly higher or lower MPRA activity under ER stress vs. DMSO (row annotation (right); red and blue bars, respectively). For each SNP (rows), z-scores are obtained by centering and scaling the normalized RNA/DNA ratios for the reference and alternate alleles (5 replicates each: DMSO and TG). Annotation horizontal bars on the left indicate whether an element (row) harbors a caQTL or T2D-associated SNP. Note that the majority of elements with higher MPRA activity under ER stress contain T2D-associated SNPs. **(b)** SNPs with allelic skew (any condition) are categorized as having at least 1 allele with – i) lower, ii) no difference in, or iii) higher MPRA activity under ER stress vs. DMSO (x-axis). SNPs in each of the above categories are further fractioned as being – i) caQTLs (red), control SNPs (yellow), or iii) T2D-associated (green). A significant proportion of caQTL SNPs with allelic skew were significantly more likely to show reduced MPRA activity, and significantly less likely to show higher MPRA activity, under ER stress. A significant proportion of T2D-associated SNPs with allelic skew displayed the opposite trend: they were significantly less likely to show reduced MPRA activity, and significantly more likely to show higher MPRA activity (86/220), under ER stress. **(c)** Fraction of T2D-associated elements or 10,000 random genomic regions overlapping SINE repeats in the human genome. The T2D-associated elements are categorized based on whether the elements show – i) MPRA activity, ii) Allelic skew in MPRA activity, or iii) higher MPRA activity under ER stress. A significant fraction of T2D-associated elements with higher activity under ER stress overlap SINEs. **(d)** The probability that elements containing SNPs with allelic skew are conserved with sequence similarity greater than 20% (y-axis) is plotted for 20 mammals whose genomes are available (x-axis). Red and yellow dots show comparisons for caQTL and control SNPs, respectively. T2D-associated SNPs with allelic skew were split based on whether they had significantly higher MPRA activity under ER stress (n=86; purple), or no change (n=110; green). Compared to 10,000 random 200 bps regions from the human genome, elements overlapping caQTL and control SNPs were significantly more likely, and T2D-associated elements with higher MPRA activity under ER stress significantly less likely, to be conserved in non-primate mammalian species (Fisher’s exact test; FDR<5%). For Panels B and C, Fisher’s Exact Test p-value (corrected for multiple testing) < 0.001 is indicated as ***, ** indicates p-value < 0.01, and * indicates p-value < 0.05.

We previously characterized islet caQTLs from individuals with intact insulin expression and secretion, and observed an enrichment of islet-specific TF binding motifs at caQTL regulatory elements^11^. ER stress, however, leads to the inactivation of β cell specific TFs (Figure 2d), with a concomitant decrease in *Ins2* gene expression (Figure 2b). As might be expected based on these observations, steady state islet caQTL-containing sequences predominantly exhibited lower MPRA activity in TG-treated cells than the DMSO solvent controls. (Figure 7a, red dashes and 7b, red). In contrast, T2D SNP-containing elements were significantly more likely to have higher MPRA activity under ER stress (Figure 7a, green and 7b). Given the majority of SNPs tested in this set are non-causal/functional ‘passenger’ alleles, this effect may reflect general CRE architecture across T2D-associated loci.

Curiously, only 13% (30/220) of T2D-associated SNPs with allelic skew in MPRA activity overlapped islet ATAC-seq peaks. We hypothesized that this may partially be due to mappability issues since 63.1% (139 / 220) of T2D-associated SNPs with allelic skew in MPRA activity overlapped repetitive sequences. Since there are no mappability issues with MPRA, we first asked if repetitive sequences have MPRA activity. Among the various classes of repetitive sequences, only elements overlapping short interspersed nuclear elements (SINEs) were significantly more likely to be active (Supplementary Figure 6a). Since >50% of elements overlapping SINEs in the MPRA library harbored T2D-associated SNPs (Supplementary Figure 6b and S6c), we next asked whether T2D-associated elements overlapping SINEs showed higher activity under ER stress. Indeed, >75% of T2D-associated elements with higher MPRA activity under ER stress overlapped SINEs, a significantly higher proportion compared to T2D-associated elements with lower or no change in MPRA activity under ER stress (Figure 7c). *Alu* elements, the most common SINEs in the human genome, are primate-specific^49^. When we investigated conservation in 20 mammalian genomes, T2D-associated elements with higher MPRA activity under ER stress were indeed less likely to be conserved in all non-primate mammalian species for which comparisons were made (Figure 7d).

In conclusion, we have identified 220 elements whose MPRA activity is disrupted by T2D-associated SNPs. 40% (n=86) of these T2D-associated SNPs had significantly higher MPRA activity under ER stress, >75% of which overlapped SINEs.

## Discussion

In this study, we used MPRA to test >13,000 sequences containing each allele of 6628 SNPs for MPRA activity in MIN6 β cells under standard culture conditions and after ER stress or paired solvent control exposures. In total, 30% (n=1983/6628) of putative CREs tested increased transcriptional activity of a minimal promoter. SNP alleles, including those associated with altered *in vivo* chromatin accessibility in human islets and with T2D genetic risk by GWAS, altered MPRA activity for approximately half of these elements (n=879/1983).

MPRA activity exhibited a striking, positive correlation with *in vivo* islet chromatin accessibility. As anticipated, elements accessible *in vivo* were far more likely to have MPRA activity *in vitro*. However, MPRA refined our understanding of the specific content and location of DNA sequences within ATAC-seq peaks that drive β cell transcriptional activation. Likelihood of an element showing MPRA activity increased as a function of its proximity to ATAC-seq peak summits, and SNPs closer to ATAC-seq peak summits were more likely to alter MPRA activity and chromatin accessibility. These results help to prioritize sequences within open chromatin regions for their importance in regulating enhancer activity by disrupting TF binding. Interactions between neighboring SNPs sometimes led to discordant effects between chromatin accessibility and activity measured with MPRA. In these instances, MPRA helped to determine the relative contributions of neighboring SNPs (e.g., additive or antagonistic effects) on transcriptional activation.

T2D-associated SNPs are overwhelmingly non-coding, and primarily alter regulatory activity rather than protein structure and function. A key challenge in T2D genetics is to identify the putative causal SNP(s) from among tens to hundreds of genetically linked, noncoding variants per association signal. This is the first study to systematically test thousands of T2D-associated SNP alleles for their effects on transcriptional activity in β cells. MPRA identified 492 elements in T2D-associated loci as active. 220/492 SNP alleles, representing 104 distinct T2D association signals, significantly altered β cell MPRA activity. MPRA recapitulated, and thereby confirmed, allelic effects of several T2D SNPs that have been characterized previously using targeted, low throughput luciferase assays, such as rs10428126 and rs2943656 at the *IGF2BP2* and *IRS1* locus, respectively. For 54/104 T2D association signals, only one SNP showed an allelic skew among all the SNPs tested, implicating them as the putative causal SNP within their respective locus. Although not an exhaustive test of all credible set SNPs identified to date, this approach has substantially expanded the list of T2D-associated SNP alleles that empirically alter ß cell regulatory sequence activity. These results should facilitate targeted follow-up experiments and integrated genomic analyses for a better understanding of the functional genetics of islet (dys)function and T2D risk and strongly motivate future studies of more comprehensive panels of T2D-associated SNPs for their transcription-modulating effects in beta cells and other diabetes-relevant cell types under steady state, stimulatory, and/or stress conditions.

Islet dysfunction and beta cell failure in T2D results from both genetic and environmental risk factors. MPRA of sequences containing T2D-associated SNPs, when tested under control and ER stress conditions, revealed the effects of these factors on ß cell transcriptional activation. A significant proportion (40%; 86/220) of sequences containing T2D-associated index or linked SNPs exhibiting allelic skew under control conditions had significantly higher MPRA activity under ER stress, suggesting – i) MIN6 cells cultured under standard, high glucose (25 mM) conditions are already under a basal level of ER stress, and ii) many T2D-associated SNPs overlap CREs relevant for β cells to respond to ER stress. Uncompensated ER stress was found to lead to the inactivation of β cell-specific TFs causing the downregulation of insulin transcription and secretion^50–54^. ER folding and trafficking capacity may therefore be a major factor determining how much insulin can be released by β cells under elevated blood glucose levels, before stress ensues. SNPs at regulatory elements modulating the expression of genes relevant to meet higher demands for insulin synthesis may impair or improve ER capacity, ultimately determining the threshold at which ER stress ensues, leading to β cell failure.

Finally, we uncovered evidence that repetitive element sequences, most notably SINEs, elicited robust transcriptional activation in ß cells and that multiple T2D-associated SNPs residing in these sequences modulated their transcriptional activity. The MPRA results obtained in this study thus suggest that repetitive element-containing sequences may play important roles in modulating stimulus and/or stress-responsive ß cell transcriptional programs and remind us to consider the potentially important roles that repetitive elements, and SNPs within them, may play in the genetics of islet (dys)function, diabetes risk and progression. Studies over the past few years demonstrating repetitive element-mediated oncogene activation and modulation of chromatin structure have contributed to an emerging appreciation of the importance of these sequences in epigenetic and transcriptional regulation^55–63^. Recently, Hernandez et al found that Alu elements are transcriptionally induced by cellular stress, including thermal and ER stress, and that the corresponding SINE RNAs function as critical transcriptional switches during stress^64^. Future studies to elucidate the target genes of these and other MPRA active, SINE-containing regulatory sequences, will be necessary to fully understand the functional consequences of sequence variation in these transcriptionally active sequences and the potential role(s) of exaptation^57, 65, 66^ in the genetics of islet (dys)function and T2D.

## Supplementary Figure Legends

**Supplementary Figure 1: The chromatin accessibility profile of MIN6 mouse β cells most resembles that of human islets. (a)** ATAC-seq peaks from 9 human tissues were mapped to the mouse genome (mm9) using the UCSC Genome Browser liftover tool and compared to MIN6 ATAC-seq peak locations. **(b)** Heatmap showing the enrichment of islet-specific TF motifs at human ATAC-seq peaks uniquely overlapping MIN6 ATAC-seq peaks.

**Supplementary Figure 2: QC Metrics of MPRA libraries. (a)** Scatter plot of the first 2 principal components, which together explain >99% of the variation in 5 plasmid and 5 RNA MPRA replicates. RNA replicates were obtained after transfection of the MPRA library into MIN6 cells under standard culture conditions. **(b)** Pairwise Pearson correlation coefficients are shown as a heatmap with unsupervised row and column clustering.

**Supplementary Figure 3: Human-Mouse sequence similarity of elements with MPRA activity.** Human-mouse sequence similarity for elements with (red) or without (black) MPRA activity is plotted. Human elements with 0% sequence similarity did not liftover to the mouse genome (mm9) with even 1% sequence similarity. A bootstrap hypothesis test of equality did not find a statistically significant difference between elements with (red) or without (black) MPRA activity for human-mouse sequence similarity. The upper and lower end-points for equality is indicated as a blue reference band.

**Supplementary Figure 4: Enrichment of islet-specific TF binding motifs at islet ATAC-seq peak summits.** Islet ATAC-seq peak summits are significantly enriched for many islet-specific TF motifs compared to genomic regions that overlap ATAC-seq peaks but are further away from the summit.

**Supplementary Figure 5: Allelic skew in MPRA activity for 879 SNPs across the 3 experimental conditions.** (Top) Venn diagrams showing the number of SNPs with allelic skew in MPRA activity (FDR<10%) in each condition. (Bottom) Scatter plot of log fold change in MPRA activity for SNPs with allelic skew in 2 experimental conditions. SNPs for which comparisons are made are indicated as pink in the corresponding Venn diagram above each scatter plot.

**Supplementary Figure 6: The majority of elements overlapping SINEs harbor T2D-associated SNPs. (a)** Barplot showing the odds of MPRA active sequences (at least one allele; any experimental condition) overlapping long interspersed nuclear element (LINE), long terminal repeat (LTR), or short interspersed nuclear element (SINE) repetitive element classes. ***p< 0.001, Fisher’s Exact test. **(b)** Fraction of elements containing i) caQTL, ii) control, or iii) T2D-associated SNPs that overlap SINEs. 10,000 random genomic regions from hg19 are included for context. **(c)** Stacked barplot showing the fraction of elements in the MPRA library overlapping SINEs in the human genome.

## Methods

### MPRA library design

200 base pair sequences, with 100 bps flanking each side of 6628 SNPs were included in our MPRA library. The SNPs belong to 3 categories:

1. Islet chromatin accessibility quantitative trait loci (caQTLs): These are SNPs we previously identified as having a significant association with altered *in vivo* chromatin accessibility in islet samples^11^. For 1806 caQTLs, only the lead SNP was included in the MPRA library. For 94 caQTLs, the 200-bp sequences contained two SNPs in LD less than 25 bp apart. For these sequences, all four allelic combinations were synthesized and tested.
2. Control SNPs: SNPs that overlapped islet ATAC-seq peaks but did not significantly alter accessibility of those peaks were also synthesized and tested^11^. Since islet caQTLs were identified in a relatively small cohort of individuals (n=19), the following criteria were used to include SNPs for which the caQTL study was likely more powered to detect associations with chromatin accessibility:

a. unadjusted p value > 0.2,
b. minor allele frequency > 0.125 SNPs overlapping individual-specific peaks or sharing peaks with other SNPs were further filtered out. Of the remaining 15,178 SNPs, 2218 were randomly selected to be included in the MPRA library.
3. T2D-associated SNPs/indels: 200 bp sequences overlapping SNPs (n=2299), small insertions (n=72) and small deletions (n=129) in linkage disequilibrium (r^2^>0.8) with T2D-associated index SNPs (n=259) from the NHGRI/EBI GWAS Catalog (accessed January 19th, 2017) were synthesized and tested, as previously described^67^. Briefly, T2D-associated GWAS SNPs were pruned using PLINK version 1.9^68^ to identify SNPs in high linkage disequilibrium (r^2^>0.8).

The vast majority of SNPs and sequences tested belonged to only 1 of the 3 categories. However, T2D-associated SNPs overlapping 13 ATAC-seq peaks were significantly associated with chromatin accessibility in islets (caQTLs). Therefore, for analysis purposes, whenever SNPs were required to belong to only 1 of the 3 categories above (such as Figures 1A, 7A and 7B), they were not categorized as caQTLs, but as being T2D-associated only.

### MPRA library construction

The MPRA library was constructed as previously described^13^. Briefly, oligos were synthesized (Agilent Technologies) as 230 bp sequences containing 200 bp of genomic sequences and 15 bp of adaptor sequence on either end. Unique 20 bp barcodes were added by PCR along with additional constant sequence for subsequent incorporation into a backbone vector by Gibson assembly. The oligo library was expanded by electroporation into *E. coli*, and the resulting plasmid library was sequenced by Illumina 2 × 150 bp chemistry to acquire oligo-barcode pairings. The library underwent restriction digestion, and GFP with a minimal TATA promoter was inserted by Gibson assembly resulting in the 200 bp oligo sequence positioned directly upstream of the promoter and the 20 bp barcode falling in the 3’ UTR of GFP. After expansion within *E. coli* the final MPRA plasmid library was sequenced by Illumina 1 × 31 bp chemistry to acquire a baseline representation of each oligo-barcode pair within the library. Barcodes mapping to more than 1 sequence were discarded from all downstream analyses. <u>Note</u>: 2 separate batches of the MPRA library were prepared. The first batch was used to perform MPRA under standard culture conditions. This MPRA library was then electroporated into *E. coli* to obtain a second batch of the MRPA library, which was used for the DMSO-TG experiments.

### MPRA library transfection into MIN6 cells

10 million MIN6 cells were seeded in each of seven 15 cm^2^ dishes. The cells were 60-70% confluent the next day. Each 15 cm^2^ dish was replaced with 20 ml of fresh media, and transfected with 7ug of the MPRA plasmid library using 55ul Lipofectamine 2000 (38% transfection efficiency). Six hours after transfection, media was either i) not changed (MPRA under standard culture conditions), ii) replaced with media containing 250 nM Thapsigargin dissolved in 0.025% DMSO, or iii) replaced with media containing 0.025% DMSO. Thirty hours after transfection, cells were trypsinized and collected by centrifugation. Cell pellets were frozen at −80°C. For each condition (standard culture / DMSO / 250 nM TG), MIN6 cells were transfected on five separate days to generate biological replicates.

### RNA isolation and MPRA RNA-seq library generation

RNA was extracted from frozen cell pellets using the Qiagen RNeasy Midi kit. Following DNase treatment, a mixture of 3 GFP-specific biotinylated primers (Supplementary Table 1; #120, #123 and #126) were used to immunoprecipitated GFP transcripts using Streptavidin C1 Dynabeads (Life Technologies). Following another round of DNase treatment, cDNA was synthesized from GFP mRNA using SuperScript IV and purified with AMPure XP beads. Quantitative PCR using primers specific for GFP (Supplementary Table 1; #34 and #52) was used to determine the cycle at which linear amplification begins for each replicate. Replicates were diluted to approximately the same concentration based on the qPCR results, and PCR with primers #34 and #52 was used to amplify barcodes associated with the ∼13.5k sequences included in the MRPA library for each replicate (9 cycles for standard culture, and 13 cycles for DMSO / 250 nM TG). A second round of PCR (6 cycles) was used to add Illumina sequencing adaptors to the DNA/RNA replicates. The resulting MPRA barcode libraries were spiked with 5% PhiX and sequenced using Illumina single-end 31 bp chemistry (with 8 bp index read), clustered at 80-90% maximum density.

### MPRA data analysis

Data from the MPRA was analyzed as previously described^13^. Briefly, the sum of the barcode counts for each oligo within replicates was median normalized, and oligos showing differential expression relative to the plasmid input were identified by modeling a negative binomial distribution with DESeq2 and applying a false discovery rate (FDR) threshold of 1%. For sequences that displayed significant MPRA activity, a paired t-test was applied on the log-transformed RNA/plasmid ratios for each experimental replicate to test whether the reference and alternate allele had similar activity. An FDR threshold of 10% was used to identify SNPs with a significant skew in MPRA activity between alleles (allelic skew). <u>Because the</u> MPRA testing standard culture conditions was performed with a separate MRPA library preparation. Therefore, the DMSO-TG MPRA results were not directly compared to MPRA performed under standard culture conditions.

### Annotating repetitive elements tested with MPRA

The ‘RepeatMasker’ track for hg19 was downloaded from the UCSC genome browser. Among the ten different classes of repeats, only three classes (long interspersed nuclear element (LINE), long terminal repeat (LTR), and short interspersed nuclear element (SINE)) overlapped more than 100 elements tested with MPRA. Therefore, only these three classes of repeats were assessed for associations with MRPA activity.

### Mapping human regulatory sequences tested with MPRA to mammalian genomes

Liftover tool in the UCSC genome browser was used to map human sequences (hg19) tested with MPRA to 20 mammalian genomes (with a minimum ratio of 0.20 bases that must remap; allowing for multiple output regions). The 20 mammalian genomes are: papAnu2 (Baboon), felCat5 (Cat), PanTro6 (Chimpanzee), BosTau7 (Cow), canFAM3 (Dog), loxAfr3 (Elephant), nomLeu3 (Gibbon), gorGor3 (Gorilla), equCab2 (Horse), mm9 (mouse), ponAbe2 (Orangutan), aiMel1 (Panda), susScr11 (Pig), ochPri3 (Pika), oryCun2 (Rabbit), rn5 (Rat), rheMac8 (Rhesus), oviAri3 (Sheep), sorAra2 (Shrew), speTri2 (Squirrel). Human sequences that did not liftover to the genome assembly of a given species were subsequently classified as not conserved (with a minimum ratio of 0.20 bases that must remap; allowing for multiple output regions).

To obtain human-mouse sequence similarity measures, liftover was performed 99 times with the minimum ratio of bases that must remap ranging from 0.01 to 1.00 in increments of 0.01 (allowing for multiple output regions). The R package ‘sm’ was used plot density of human-mouse sequence similarity and perform non-parametric bootstrap hypothesis tests of equality. Human sequences that did not liftover to the mm9 mouse genome with even 1% sequence similarity were classified as having 0% sequence similarity.

### Cross-species mapping of human ATAC-seq peaks to MIN6 ATAC-seq peaks

Human and mouse ATAC-seq data were processed as previously described^11^. Briefly, low quality portions of reads were trimmed using Trimmomatic^70^ and aligned to the hg19 or mm9 genome assembly using Burrows Wheeler Aligner-MEM. For each replicate, duplicate reads were removed after shifting. Technical replicates were merged using SAMtools and peaks were called using MACS2^6^ (with parameters -callpeak --nomodel -f BAMPE) at FDR<1%. ATAC-seq peak summit positions were obtained from MACS2 output files. The liftover tool in the UCSC genome browser was used to map human ATAC-seq peaks to the mouse (mm9) genome using a minimum ratio of 0.10 bases that must remap (not allowing for multiple output regions). Using bedtools, human ATAC-seq peaks mapping to the mouse genome were then overlapped with MIN6 ATAC-seq peaks.

### Transcription factor (TF) motif enrichment

Homer findMotifsGenome.pl script was used to investigate TF motifs enriched in a given set of sequences. Elements with lower MPRA activity under ER stress were used as background to identify TF motifs enriched in elements with higher MPRA activity under ER stress, and vice-versa (parameters: hg19, -size given). 2,008 T2D-associated elements with no MPRA activity were used as background to identify TF motifs enriched in the 492 T2D SNP-containing sequences with significant MPRA activity (parameters: hg19, -size given). For the cross-species analysis of ATAC-seq peaks, ATAC-seq peaks shared with other human cell types were used as background to identify TF motifs enriched in unique ATAC-seq peaks (parameters: mm9, -size given).

### Analysis of islet ChIP-seq data

Chromatin immunoprecipitation sequencing (ChIP-seq) data from Pasquali et al.^17^ were aligned to the hg19 reference human genome as previously described^44^. Elements tested with MPRA were then overlapped with ChIP-seq peaks to conduct Fisher’s exact tests using R.

### CentriMo analysis of ATAC-seq peak summits

CentriMo software in the MEME Suite was used to identify TF motifs enriched in the 100 bp genomic regions flanking human islet ATAC-seq peak summits^71^. Negative controls were 200 bp genomic regions overlapping ATAC-seq peaks but flanking the human islet ATAC-seq peak summits (default parameters).

## Acknowledgements

We thank members of the Stitzel and Ucar labs for helpful discussion and critiques during study design and execution and Francis S. Collins, D. Leland Taylor, Christine Beck and Taneli Helenius for helpful comments on the manuscript. This work was supported by W81XWH-18-0401 (to MLS and DU) and R00HG008179 (to RT).

---

| IndexSNP | Locus | SNP | REF_ALLELE | ALT_ALLELE | MAF_AFR | MAF_AMR |
| --- | --- | --- | --- | --- | --- | --- |
| rs7903146 | TCF7L2 | rs7903146 | C | T | 0.28 | 0.25 |
| rs9472138 | VEGFA | rs12189774 | G | A | 0.19 | 0.21 |
| rs3842770 | INS-IGF2 | rs115420895 | G | A | 0.06 | 0.01 |
| rs7578326 | IRS1 | rs2943656 | A | G | 0.6 | 0.73 |
| rs10507349 | RNF6 | rs4630391 | C | T | 0.08 | 0.36 |
| rs1535500 | KCNK16 | rs12663159 | C | A | 0.86 | 0.49 |
| rs864745 | JAZF1 | rs1635852 | T | C | 0.22 | 0.4 |
| rs864745 |  | rs849133 | C | T | 0.21 | 0.4 |
|  |  | rs10811660 | G | A | 0.05 | 0.14 |
| rs10811661 | CDKN2B-AS1 | rs10811661 | T | C | 0.05 | 0.14 |
|  |  | rs10965246 | T | C | 0.07 | 0.14 |
|  |  | rs10428126 | T | C | 0.6 | 0.29 |
|  |  | rs9808924 | G | A | 0.8 | 0.32 |
| rs1470579 | IGF2BP2 | rs9854769 | A | G | 0.8 | 0.31 |
|  |  | rs7630554 | A | G | 0.79 | 0.31 |
|  |  | rs35188816 | CA | C | 0.78 | 0.31 |
| rs1048886 | SDHAF4 | rs2881632 | C | A | 0.4 | 0.22 |
| rs12496230 | KBTBD8 | rs13096599 | T | C | 0.02 | 0.2 |
| rs6467136 | GCC1 | rs7783500 | G | A | 0.54 | 0.4 |
| rs10923931 | NOTCH2 | rs72697237 | A | T | 0.36 | 0.12 |
| rs7578597 | THADA | rs13405776 | C | T | 0.34 | 0.07 |
| rs7630877 | PEX5L | rs17748864 | C | T | 0.24 | 0.27 |
| rs7957197 | OASL | rs7957197 | T | A | 0.17 | 0.15 |
| rs34669198 | SRBD1 | rs13026123 | T | A | 0.09 | 0.09 |

| Allele with higher MPRA activity under |  |  |  |  |
| --- | --- | --- | --- | --- |
| MAF_ASN | MAF_EUR | Standard Culture | DMSO | 250 nM TG |
| 0.03 | 0.31 | - | T | - |
| 0.11 | 0.29 | - | G | G |
| 0 | 0.0013 | A | - | - |
| 0.94 | 0.62 | - | G | G |
| 0.44 | 0.24 | C | C | C |
| 0.43 | 0.49 | A | - | - |
| 0.21 | 0.54 | - | C | - |
| 0.21 | 0.53 | C | C | - |
| 0.44 | 0.16 | - | G | - |
| 0.44 | 0.16 | - | C | - |
| 0.44 | 0.17 | C | C | - |
| 0.28 | 0.29 | T | T | - |
| 0.29 | 0.29 | A | - | - |
| 0.29 | 0.29 | G | G | G |
| 0.29 | 0.28 | G | G | G |
| 0.28 | 0.27 | - | CA | - |
| 0.12 | 0.19 | A | A | A |
| 0.08 | 0.18 | C | C | C |
| 0.67 | 0.42 | G | G | G |
| 0.03 | 0.11 | T | T | T |
| 0.01 | 0.09 | T | T | T |
| 0.15 | 0.34 | T | T | T |
| 0 | 0.2 | T | T | T |
| 0.1 | 0.13 | T | T | T |

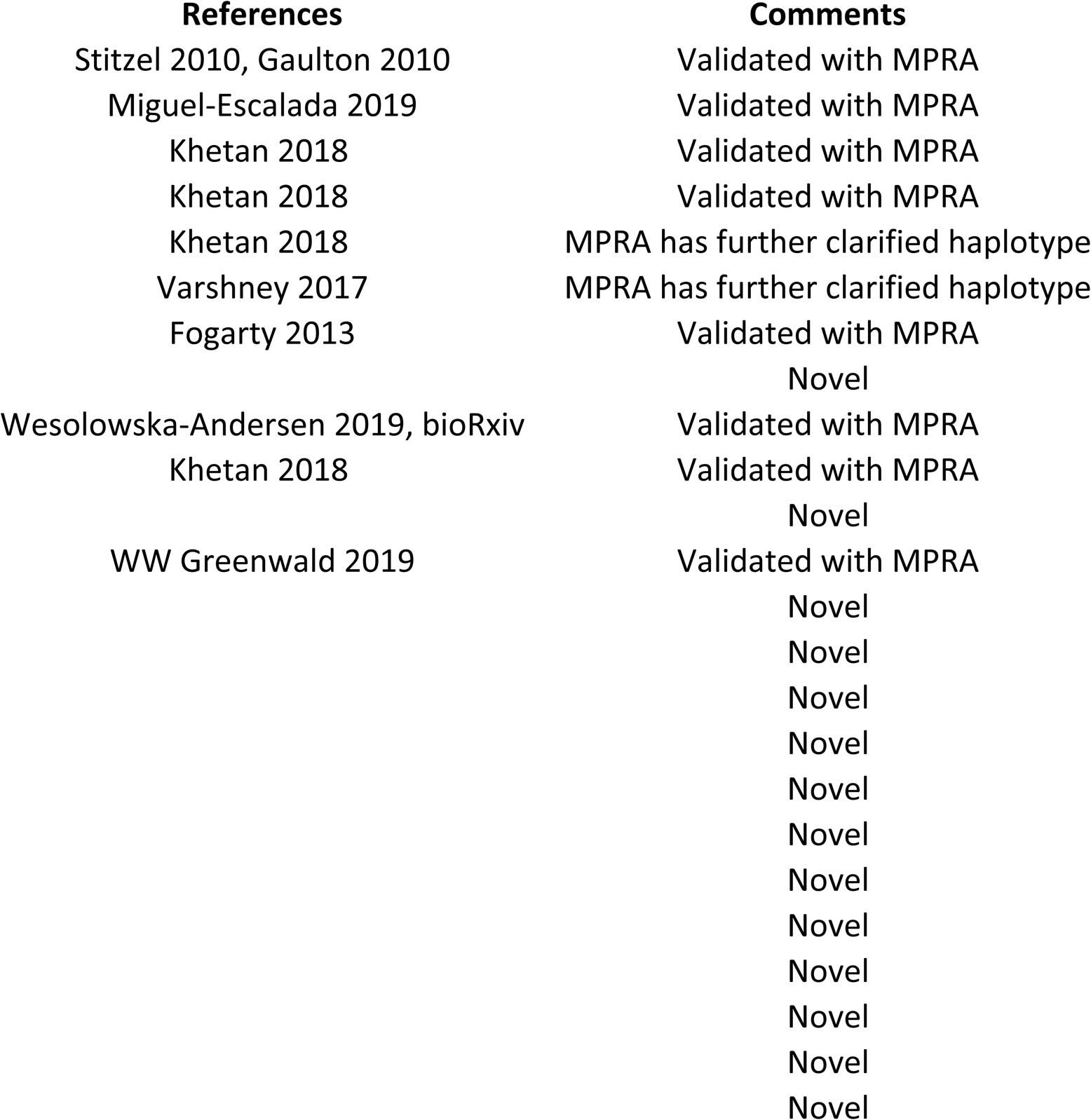

## References

1. 1000 Genomes Project Consortium et al. A global reference for human genetic variation. Nature 526, 68–74 (2015).

2. MacArthur, J. et al. The new NHGRI-EBI Catalog of published genome-wide association studies (GWAS Catalog). Nucleic Acids Res. 45, D896–D901 (2017).

3. Mahajan, A. et al. Fine-mapping of an expanded set of type 2 diabetes loci to single-variant resolution using high-density imputation and islet-specific epigenome maps. (2018) doi:10.1101/245506.

4. Fuchsberger, C. et al. The genetic architecture of type 2 diabetes. Nature 536, 41–47 (2016).

5. Buenrostro, J. D., Giresi, P. G., Zaba, L. C., Chang, H. Y. & Greenleaf, W. J. Transposition of native chromatin for fast and sensitive epigenomic profiling of open chromatin, DNA-binding proteins and nucleosome position. Nat Meth 10, 1213–1218 (2013).

6. Zhang, Y. et al. Model-based analysis of ChIP-Seq (MACS). Genome Biol. 9, R137 (2008).

7. Wang, X. et al. High-resolution genome-wide functional dissection of transcriptional regulatory regions and nucleotides in human. Nat. Commun. 9, 5380 (2018).

8. Movva, R. et al. Deciphering regulatory DNA sequences and noncoding genetic variants using neural network models of massively parallel reporter assays. PloS One 14, e0218073 (2019).

9. Degner, J. F. et al. DNase I sensitivity QTLs are a major determinant of human expression variation. Nature 482, 390–394 (2012).

10. Kumasaka, N., Knights, A. J. & Gaffney, D. J. Fine-mapping cellular QTLs with RASQUAL and ATAC-seq. Nat. Genet. 48, 206–213 (2016).

11. Khetan, S. et al. Type 2 Diabetes-Associated Genetic Variants Regulate Chromatin Accessibility in Human Islets. Diabetes 67, 2466–2477 (2018).

12. Melnikov, A. et al. Systematic dissection and optimization of inducible enhancers in human cells using a massively parallel reporter assay. Nat. Biotechnol. 30, 271–277 (2012).

13. Tewhey, R. et al. Direct Identification of Hundreds of Expression-Modulating Variants using a Multiplexed Reporter Assay. Cell 165, 1519–1529 (2016).

14. Ulirsch, J. C. et al. Systematic Functional Dissection of Common Genetic Variation Affecting Red Blood Cell Traits. Cell 165, 1530–1545 (2016).

15. Vockley, C. M. et al. Massively parallel quantification of the regulatory effects of noncoding genetic variation in a human cohort. Genome Res. 25, 1206–1214 (2015).

16. Klein, J. C. et al. Functional testing of thousands of osteoarthritis-associated variants for regulatory activity. Nat. Commun. 10, 2434 (2019).

17. Pasquali, L. et al. Pancreatic islet enhancer clusters enriched in type 2 diabetes risk-associated variants. Nat. Genet. 46, 136–143 (2014).

18. Parker, S. C. J. et al. Chromatin stretch enhancer states drive cell-specific gene regulation and harbor human disease risk variants. Proc. Natl. Acad. Sci. U. S. A. 110, 17921–17926 (2013).

19. Roman, T. S. et al. A Type 2 Diabetes-Associated Functional Regulatory Variant in a Pancreatic Islet Enhancer at the Adcy5 Locus. Diabetes (2017) doi:10.2337/db17-0464.

20. Kycia, I. et al. A common type 2 diabetes risk variant potentiates activity of an evolutionarily conserved islet stretch enhancer and increases C2CD4A/B expression. Am. J. Hum. Genet. in press, (2018).

21. Varshney, A. et al. Genetic regulatory signatures underlying islet gene expression and type 2 diabetes. Proc. Natl. Acad. Sci. U. S. A. (2017) doi:10.1073/pnas.1621192114.

22. Sharma, R. B. et al. Insulin demand regulates β cell number via the unfolded protein response. J. Clin. Invest. 125, 3831–3846 (2015).

23. Dooley, J. et al. Genetic predisposition for beta cell fragility underlies type 1 and type 2 diabetes. Nat. Genet. 48, 519–527 (2016).

24. Bonnycastle, L. L. et al. Autosomal dominant diabetes arising from a Wolfram syndrome 1 mutation. Diabetes 62, 3943–3950 (2013).

25. Delépine, M. et al. EIF2AK3, encoding translation initiation factor 2-alpha kinase 3, is mutated in patients with Wolcott-Rallison syndrome. Nat. Genet. 25, 406– 409 (2000).

26. Booth, C. & Koch, G. L. Perturbation of cellular calcium induces secretion of luminal ER proteins. Cell 59, 729–737 (1989).

27. Sehgal, P. et al. Inhibition of the sarco/endoplasmic reticulum (ER) Ca2+-ATPase by thapsigargin analogs induces cell death via ER Ca2+ depletion and the unfolded protein response. J. Biol. Chem. 292, 19656–19673 (2017).

28. Stone, S. et al. Pancreatic stone protein/regenerating protein is a potential biomarker for endoplasmic reticulum stress in beta cells. Sci. Rep. 9, 5199 (2019).

29. Robbins, R. D. et al. Inhibition of deoxyhypusine synthase enhances islet {beta} cell function and survival in the setting of endoplasmic reticulum stress and type 2 diabetes. J. Biol. Chem. 285, 39943–39952 (2010).

30. Cunha, D. A. et al. Pancreatic β-cell protection from inflammatory stress by the endoplasmic reticulum proteins thrombospondin 1 and mesencephalic astrocyte-derived neutrotrophic factor (MANF). J. Biol. Chem. 292, 14977–14988 (2017).

31. Syed, I. et al. PAHSAs attenuate immune responses and promote β cell survival in autoimmune diabetic mice. J. Clin. Invest. 129, 3717–3731 (2019).

32. Chen, Y.-C. et al. PAM haploinsufficiency does not accelerate the development of diet- and human IAPP-induced diabetes in mice. Diabetologia 63, 561–576 (2020).

33. Suetomi, R. et al. Adrenomedullin has a cytoprotective role against ER stress for pancreatic ß-cells in autocrine and paracrine manners. J. Diabetes Investig. (2020) doi:10.1111/jdi.13218.

34. Gao, N. et al. Foxa1 and Foxa2 maintain the metabolic and secretory features of the mature beta-cell. Mol. Endocrinol. Baltim. Md 24, 1594–1604 (2010).

35. Fu, Z., Gilbert, E. R. & Liu, D. Regulation of insulin synthesis and secretion and pancreatic Beta-cell dysfunction in diabetes. Curr. Diabetes Rev. 9, 25–53 (2013).

36. Cullinan, S. B. & Diehl, J. A. Coordination of ER and oxidative stress signaling: the PERK/Nrf2 signaling pathway. Int. J. Biochem. Cell Biol. 38, 317–332 (2006).

37. Oyadomari, S. & Mori, M. Roles of CHOP/GADD153 in endoplasmic reticulum stress. Cell Death Differ. 11, 381–389 (2004).

38. Efanov, A. M., Sewing, S., Bokvist, K. & Gromada, J. Liver X receptor activation stimulates insulin secretion via modulation of glucose and lipid metabolism in pancreatic beta-cells. Diabetes 53 Suppl 3, S75–78 (2004).

39. Ogihara, T. et al. Liver X receptor agonists augment human islet function through activation of anaplerotic pathways and glycerolipid/free fatty acid cycling. J. Biol. Chem. 285, 5392–5404 (2010).

40. Aguayo-Mazzucato, C. et al. Thyroid hormone promotes postnatal rat pancreatic β-cell development and glucose-responsive insulin secretion through MAFA. Diabetes 62, 1569–1580 (2013).

41. Aguayo-Mazzucato, C. et al. T3 Induces Both Markers of Maturation and Aging in Pancreatic β-Cells. Diabetes 67, 1322–1331 (2018).

42. Brun, P.-J. et al. Retinoic acid receptor signaling is required to maintain glucose-stimulated insulin secretion and β-cell mass. FASEB J. Off. Publ. Fed. Am. Soc. Exp. Biol. 29, 671–683 (2015).

43. Peiris, H. et al. Discovering human diabetes-risk gene function with genetics and physiological assays. Nat. Commun. 9, 3855 (2018).

44. Stitzel, M. L. et al. Global epigenomic analysis of primary human pancreatic islets provides insights into type 2 diabetes susceptibility loci. Cell Metab. 12, 443–455 (2010).

45. Gaulton, K. J. et al. A map of open chromatin in human pancreatic islets. Nat. Genet. 42, 255–259 (2010).

46. Fogarty, M. P., Panhuis, T. M., Vadlamudi, S., Buchkovich, M. L. & Mohlke, K. L. Allele-specific transcriptional activity at type 2 diabetes-associated single nucleotide polymorphisms in regions of pancreatic islet open chromatin at the JAZF1 locus. Diabetes 62, 1756–1762 (2013).

47. Greenwald, W. W. et al. Pancreatic islet chromatin accessibility and conformation reveals distal enhancer networks of type 2 diabetes risk. Nat. Commun. 10, 2078 (2019).

48. Miguel-Escalada, I. et al. Human pancreatic islet three-dimensional chromatin architecture provides insights into the genetics of type 2 diabetes. Nat. Genet. 51, 1137–1148 (2019).

49. Deininger, P. Alu elements: know the SINEs. Genome Biol. 12, 236 (2011).

50. Kono, T. et al. Impaired Store-Operated Calcium Entry and STIM1 Loss Lead to Reduced Insulin Secretion and Increased Endoplasmic Reticulum Stress in the Diabetic β-Cell. Diabetes 67, 2293–2304 (2018).

51. Ghiasi, S. M. et al. Endoplasmic Reticulum Chaperone Glucose-Regulated Protein 94 Is Essential for Proinsulin Handling. Diabetes 68, 747–760 (2019).

52. Xin, Y. et al. Pseudotime Ordering of Single Human β-Cells Reveals States of Insulin Production and Unfolded Protein Response. Diabetes 67, 1783–1794 (2018).

53. Kim, M. J. et al. Attenuation of PERK enhances glucose-stimulated insulin secretion in islets. J. Endocrinol. 236, 125–136 (2018).

54. Hasnain, S. Z., Prins, J. B. & McGuckin, M. A. Oxidative and endoplasmic reticulum stress in β-cell dysfunction in diabetes. J. Mol. Endocrinol. 56, R33–54 (2016).

55. Pehrsson, E. C., Choudhary, M. N. K., Sundaram, V. & Wang, T. The epigenomic landscape of transposable elements across normal human development and anatomy. Nat. Commun. 10, 5640 (2019).

56. Su, M., Han, D., Boyd-Kirkup, J., Yu, X. & Han, J.-D. J. Evolution of Alu elements toward enhancers. Cell Rep. 7, 376–385 (2014).

57. Jang, H. S. et al. Transposable elements drive widespread expression of oncogenes in human cancers. Nat. Genet. 51, 611–617 (2019).

58. Franchini, L. F. et al. Convergent evolution of two mammalian neuronal enhancers by sequential exaptation of unrelated retroposons. Proc. Natl. Acad. Sci. U. S. A. 108, 15270–15275 (2011).

59. Choudhary, M. N. et al. Co-opted transposons help perpetuate conserved higher-order chromosomal structures. Genome Biol. 21, 16 (2020).

60. Sundaram, V. et al. Widespread contribution of transposable elements to the innovation of gene regulatory networks. Genome Res. 24, 1963–1976 (2014).

61. Xie, M. et al. DNA hypomethylation within specific transposable element families associates with tissue-specific enhancer landscape. Nat. Genet. 45, 836–841 (2013).

62. Sundaram, V. et al. Functional cis-regulatory modules encoded by mouse-specific endogenous retrovirus. Nat. Commun. 8, 14550 (2017).

63. Batut, P., Dobin, A., Plessy, C., Carninci, P. & Gingeras, T. R. High-fidelity promoter profiling reveals widespread alternative promoter usage and transposon-driven developmental gene expression. Genome Res. 23, 169–180 (2013).

64. Hernandez, A. J. et al. B2 and ALU retrotransposons are self-cleaving ribozymes whose activity is enhanced by EZH2. Proc. Natl. Acad. Sci. U. S. A. 117, 415–425 (2020).

65. Bejerano, G. et al. A distal enhancer and an ultraconserved exon are derived from a novel retroposon. Nature 441, 87–90 (2006).

66. Mita, P. & Boeke, J. D. How retrotransposons shape genome regulation. Curr. Opin. Genet. Dev. 37, 90–100 (2016).

67. Lawlor, N. et al. Multiomic Profiling Identifies cis-Regulatory Networks Underlying Human Pancreatic β Cell Identity and Function. Cell Rep. 26, 788–801.e6 (2019).

68. Purcell, S. et al. PLINK: a tool set for whole-genome association and population-based linkage analyses. Am. J. Hum. Genet. 81, 559–575 (2007).

69. Love, M. I., Huber, W. & Anders, S. Moderated estimation of fold change and dispersion for RNA-seq data with DESeq2. Genome Biol. 15, 550 (2014).

70. Bolger, A. M., Lohse, M. & Usadel, B. Trimmomatic: a flexible trimmer for Illumina sequence data. Bioinforma. Oxf. Engl. 30, 2114–2120 (2014).

71. Bailey, T. L. & Machanick, P. Inferring direct DNA binding from ChIP-seq. Nucleic Acids Res. 40, e128 (2012).

