## Supplementary figures and images for "Functional characterization of thousands of type 2 diabetes-associated and chromatin-modulating variants under steady state and endoplasmic reticulum stress"

### Supplementary Figure 1

**Supplementary Figure 1**

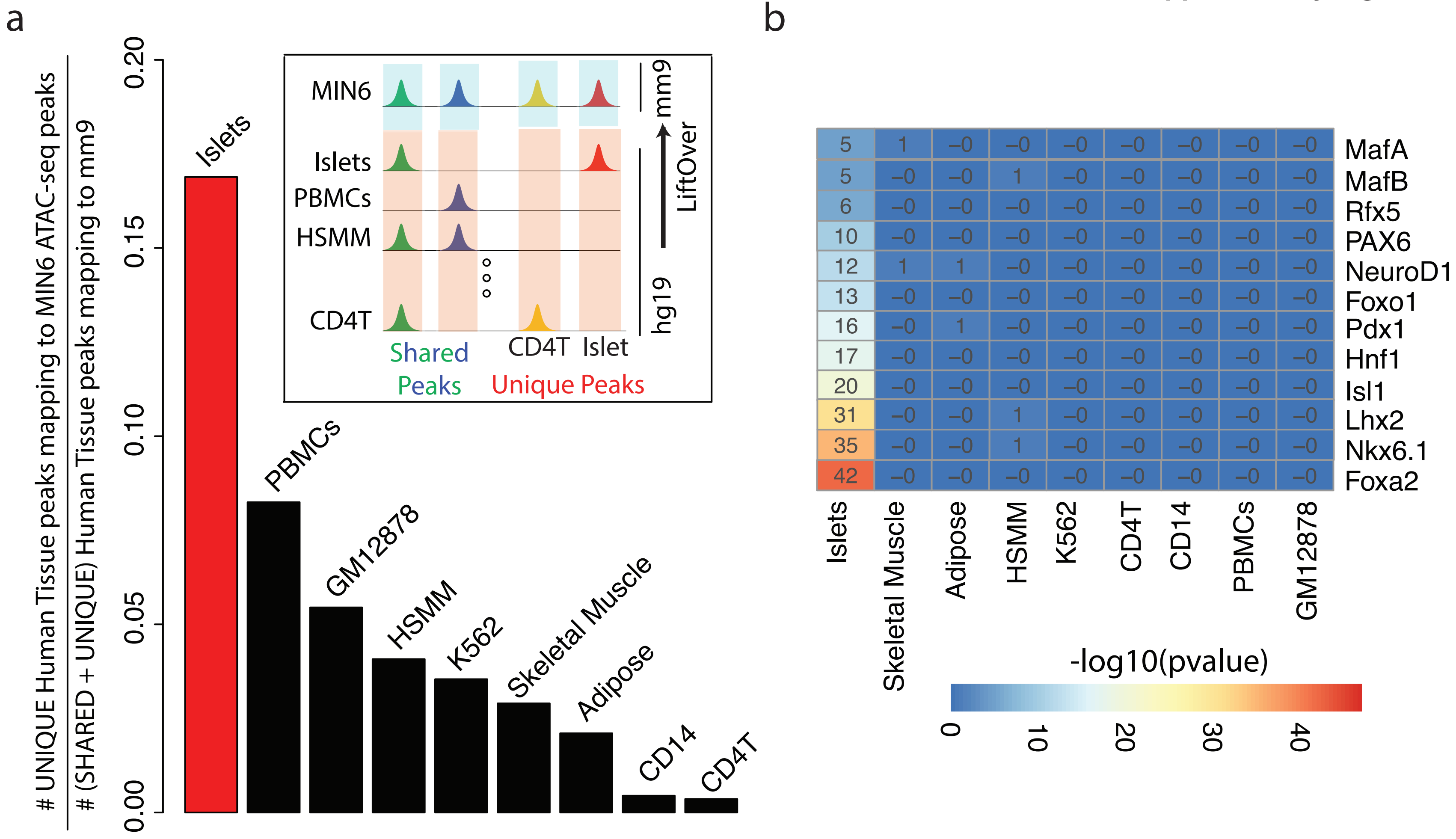

### Supplementary Figure 3

##### Supplementary Figure 3

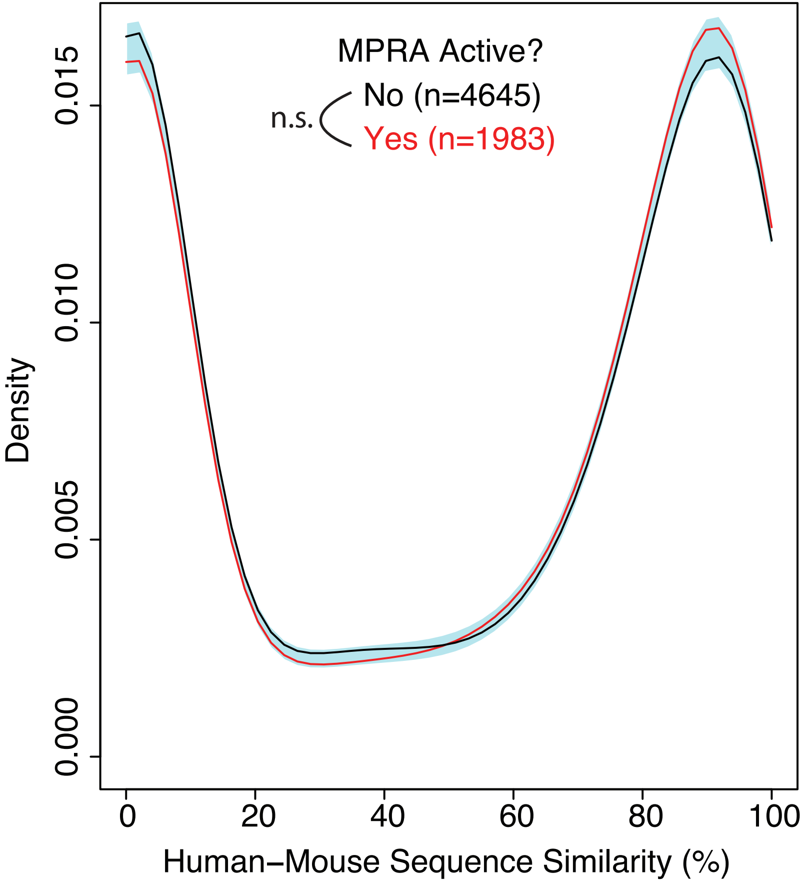

### Supplementary Figure 4

**Supplementary Figure 4**

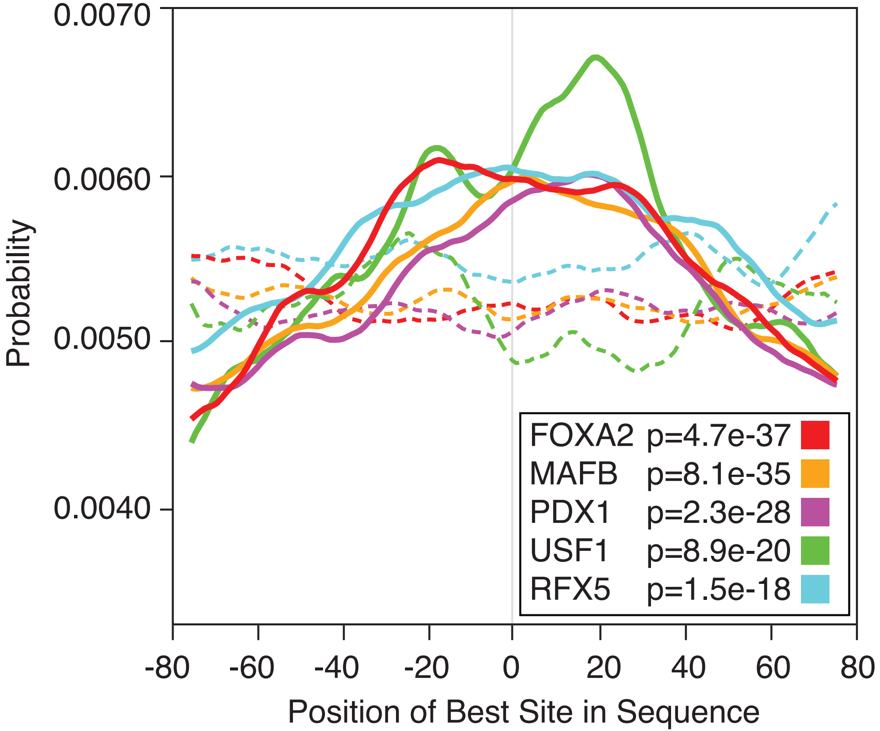

### Supplementary Figure 5

**Supplementary Figure 5**

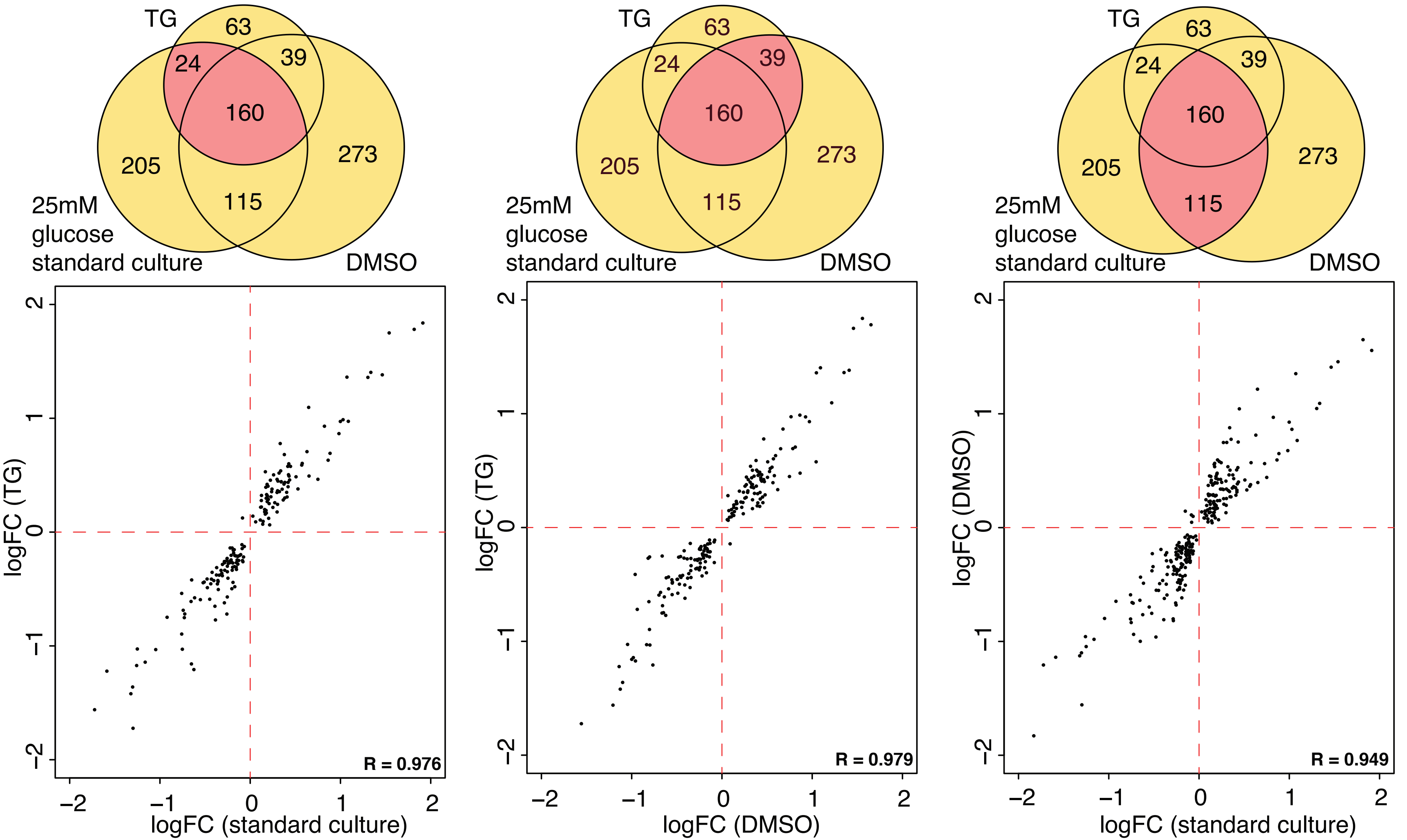

### Supplementary Figure 6

#### Supplementary Figure 6

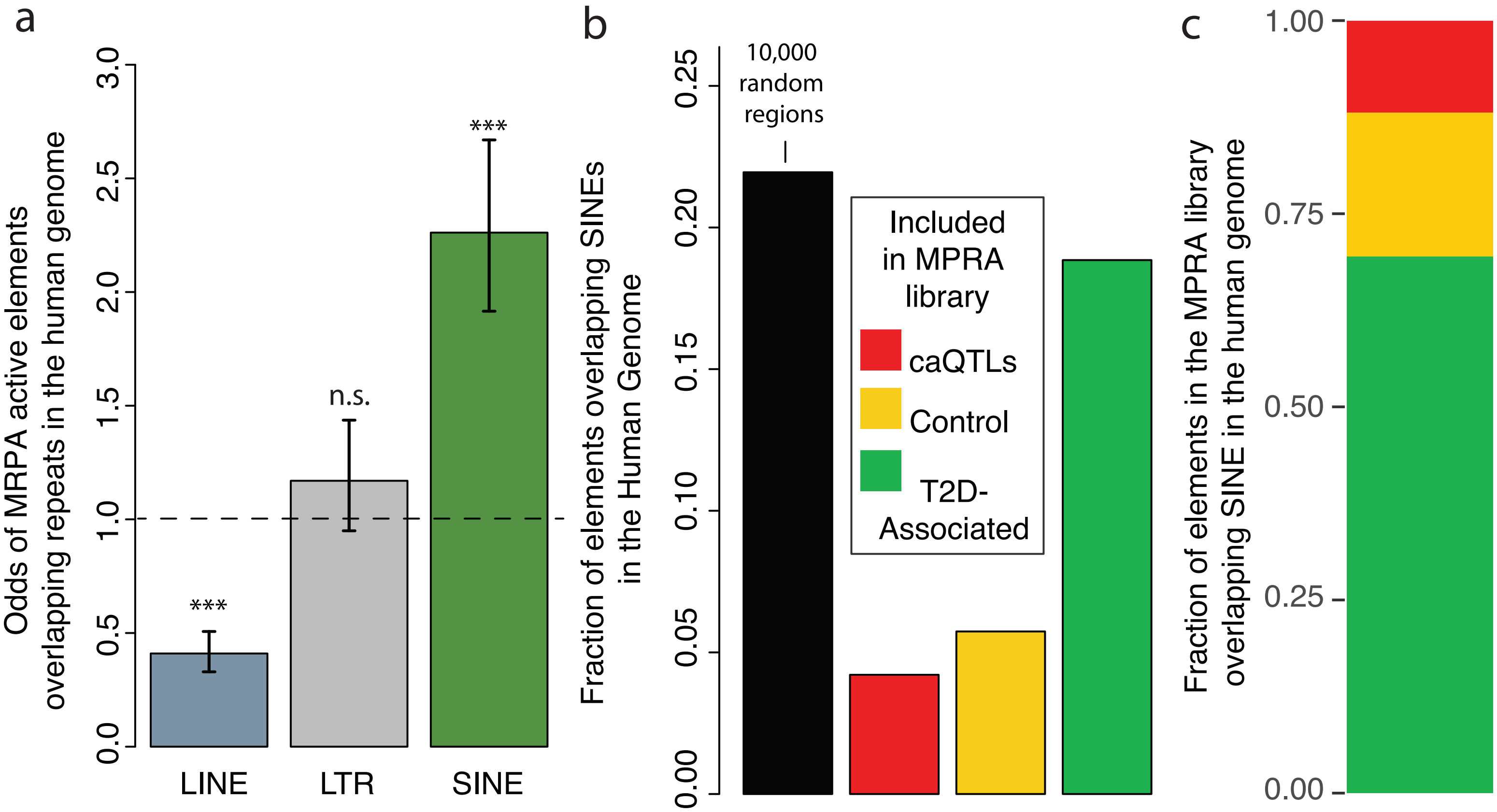
