## Supplementary Figure 2 for "Functional characterization of thousands of type 2 diabetes-associated and chromatin-modulating variants under steady state and endoplasmic reticulum stress"

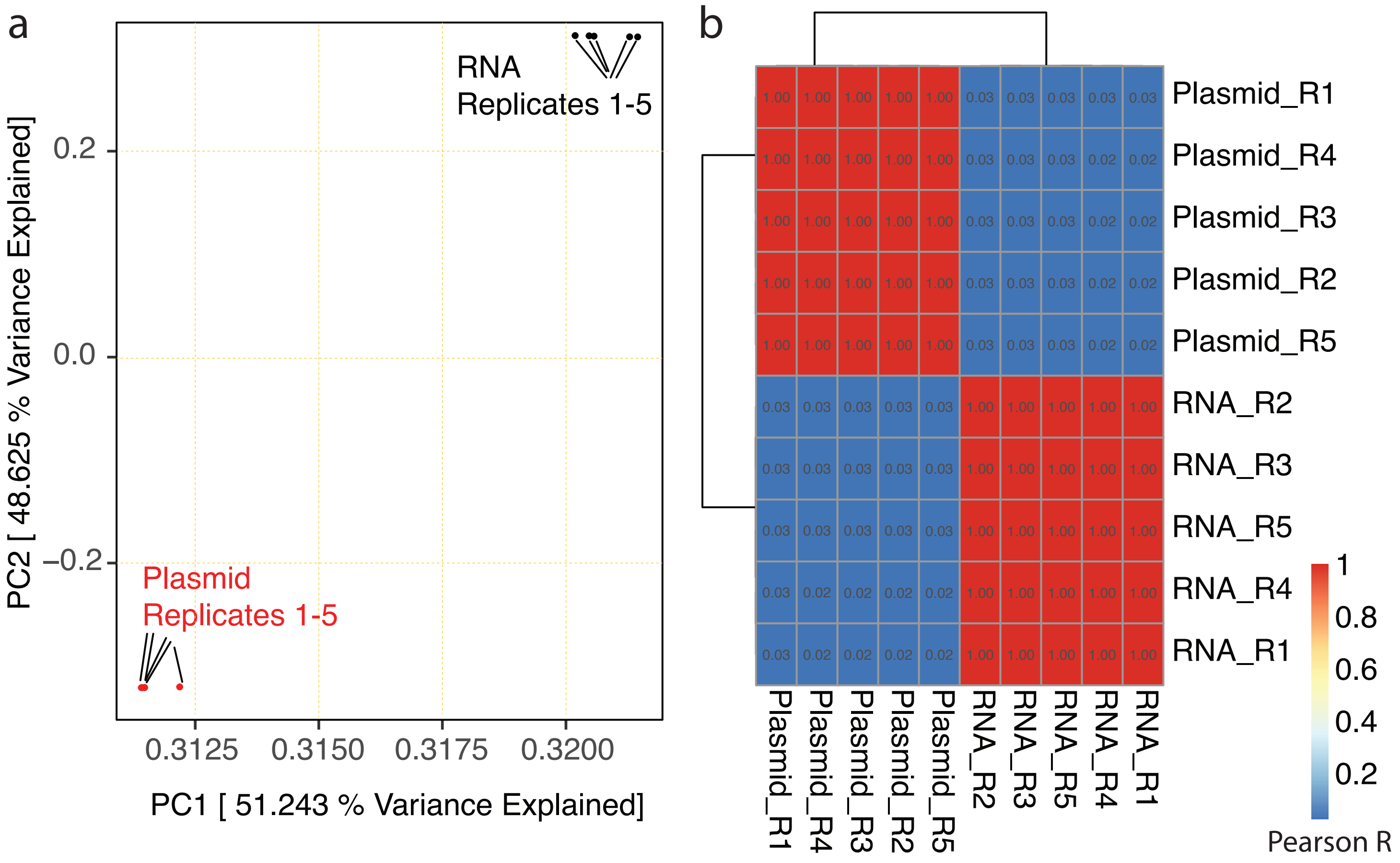


| | Plasmid_R1 | Plasmid_R4 | Plasmid_R3 | Plasmid_R2 | Plasmid_R5 | RNA_R2 | RNA_R3 | RNA_R5 | RNA_R4 | RNA_R1 |
|---|---|---|---|---|---|---|---|---|---|---|
| Plasmid_R1 | 1.00 | 1.00 | 1.00 | 1.00 | 1.00 | 0.03 | 0.03 | 0.03 | 0.03 | 0.03 |
| Plasmid_R4 | 1.00 | 1.00 | 1.00 | 1.00 | 1.00 | 0.03 | 0.03 | 0.03 | 0.02 | 0.02 |
| Plasmid_R3 | 1.00 | 1.00 | 1.00 | 1.00 | 1.00 | 0.03 | 0.03 | 0.03 | 0.02 | 0.02 |
| Plasmid_R2 | 1.00 | 1.00 | 1.00 | 1.00 | 1.00 | 0.03 | 0.03 | 0.03 | 0.02 | 0.02 |
| Plasmid_R5 | 1.00 | 1.00 | 1.00 | 1.00 | 1.00 | 0.03 | 0.03 | 0.03 | 0.02 | 0.02 |
| RNA_R2 | 0.03 | 0.03 | 0.03 | 0.03 | 0.03 | 1.00 | 1.00 | 1.00 | 1.00 | 1.00 |
| RNA_R3 | 0.03 | 0.03 | 0.03 | 0.03 | 0.03 | 1.00 | 1.00 | 1.00 | 1.00 | 1.00 |
| RNA_R5 | 0.03 | 0.03 | 0.03 | 0.03 | 0.03 | 1.00 | 1.00 | 1.00 | 1.00 | 1.00 |
| RNA_R4 | 0.03 | 0.02 | 0.02 | 0.02 | 0.02 | 1.00 | 1.00 | 1.00 | 1.00 | 1.00 |
| RNA_R1 | 0.03 | 0.02 | 0.02 | 0.02 | 0.02 | 1.00 | 1.00 | 1.00 | 1.00 | 1.00 |
